# The Effects of Heat Stress on the Ovary, Follicles and Oocytes: A Systematic Review

**DOI:** 10.1101/2024.12.04.626831

**Authors:** Luhan T. Zhou, Dilan Gokyer, Krystal Madkins, Molly Beestrum, Daniel E. Horton, Francesca E. Duncan, Elnur Babayev

## Abstract

Climate change is driving significant environmental changes with profound implications for human health, including fertility. While the detrimental effects of heat on spermatogenesis are well-documented, the impact of elevated temperatures on ovaries and female fertility remains less explored. This review systematically examines the literature on heat stress (HS) effects on mammalian ovaries, follicles, and oocytes. Evidence from mammalian models indicates that HS significantly impairs ovarian function, disrupting hormone profiles, reducing ovarian size and weight, altering histology, decreasing granulosa cell viability, and compromising oocyte quality. Efforts to develop strategies and substances to mitigate these adverse effects are ongoing, but further research into the underlying mechanisms is urgently needed.

**Summary Sentence:** This systematic review summarizes the evidence on the adverse effects of heat stress on the mammalian oocyte and ovary

## Introduction

The complex phenomena of climate change and global warming are the result of increasing greenhouse gas concentrations that have substantially altered the Earth’s environment and human vulnerability [1, 2]. Rising temperatures have resulted in altered weather patterns, more frequent and intense heatwaves, droughts, and wildfires, and in some locales worsening air quality, all leading to a global health crisis [3]. According to the World Health Organization (WHO), between 2030 and 2050, approximately 250,000 additional deaths per year will be due to climate change [4]. A report from the Red Cross found that in 2020, there were 132 unique extreme weather events with 51.6 million people having been affected globally and over 3,000 killed [5]. Overall, climate change has had and will continue to have profound implications on human health and well-being, with multifaceted impacts due to the varying geographical regions and socio-economic backgrounds of individuals. Additionally, vulnerable populations have been found to be even more susceptible to the effects of climate change [6]. This includes women, the elderly, children, pregnant women, individuals with pre-existing health conditions, and disinvested communities with limited access to resources.

Of the vulnerable populations, women and pregnant women are of particular interest as the impact of global warming and the exposure to high temperatures on fertility and reproduction is an area of increasing concern. Adverse effects of heat on spermatogenesis are well-recognized, including reduced sperm output, increased percentage of abnormal/aged sperm, reduced sperm counts and motility, and subfertility [7–9]. However, relatively less is known about the impact of elevated temperatures on female fertility and reproduction. Females are born with a finite number of oocytes, formed during fetal development, which comprise the ovarian reserve and dictates the reproductive lifespan [10]. At ∼20 weeks of gestation, the fetal ovary contains a peak of millions of primordial follicles, each consisting of an immature oocyte surrounded by supportive somatic cells [11]. As females age, the total number of follicles declines and ends at menopause with the depletion of the ovarian reserve and cessation of reproductive capacity [12, 13]. Menopause is associated with increased risk of all-cause mortality, cardiovascular disease, metabolic syndrome, musculoskeletal diseases, and neurological dysfunction [14, 15]. Delayed childbearing and reproductive aging are also associated with the increase in infertility and miscarriage rates due to decline in gamete quality and quantity [16, 17]. Any environmental process that affects the ovarian reserve, rate of its decline, and quality of gametes could have profound effects on female reproduction and overall health. Increasing average global temperatures [18] and the frequency of days with extreme temperatures [19] may negatively affect female reproduction and reproductive aging. In this manuscript, we aimed to systematically compile and closely examine the literature on the impacts of elevated temperatures on the ovary and oocyte, and review potential mechanisms of and mitigating strategies for these effects.

## Materials and Methods

We performed a systematic review, and articles were identified through a search of PubMed (NIH/NLM), Embase (Elsevier), Cochrane Database of Systematic Reviews and Central Register of Controlled Trials (Wiley), Scopus (Elsevier), and Global Health (EBSCOhost) through October 17, 2024. The search strategy was designed by academic librarians (MB and KM) and involved an extensive list of key words and database-specific controlled vocabulary related to fertility, ovaries and heat stress **(Supplementary Table S1)**. Eligible studies were limited to the English language, mammalian species, and full-text articles. Meeting abstracts, comments, case reports, news, and letters were excluded. There was no limitation on the date of publication and all citations through October 2024 were included. The review strategy followed the preferred reporting items for systematic review and meta-analysis (PRISMA) guidelines and is summarized in **Fig. 1**.

**Figure 1.**
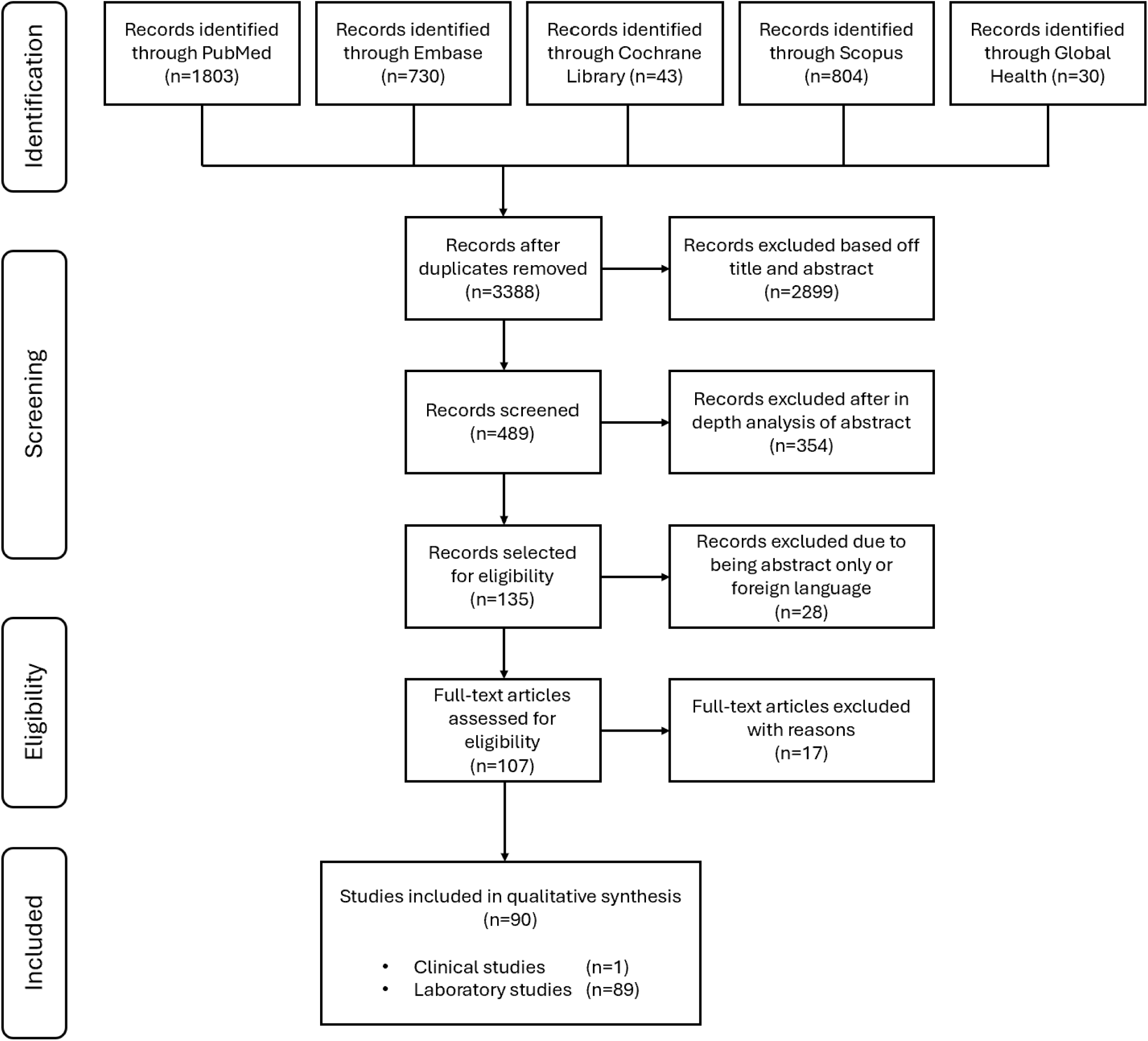
PRISMA flow chart of study selection. Ninety articles met all the inclusion criteria and were included in a qualitative synthesis. PRISMA, preferred reporting items for systematic review and meta-analysis. Seventeen studies were excluded after full text review as they either studied cooling effects, did not specify their HS paradigm, only focused on HS, or did not report controls, control temperatures, and/or the duration of the experimental intervention.

Out of the 3410 initially identified articles, 90 addressed the review scope through cross referencing using Rayyan [20] (LTZ and DG) and were fully extracted in the tabular form. Of the 90 included studies, 89 were laboratory studies and 1 was a clinical study **(Fig. 1)**. This review only included the studies that investigated the effects of increased temperature on the ovary, follicle, oocyte, and/or granulosa cells, where both control and experimental groups were analyzed. Seventeen studies were excluded after full text review as they either studied cooling effects, did not specify their heat stress (HS) paradigm, or did not report controls, control temperatures, and/or the duration of the experimental intervention [21–37].

## Results

### The effects of heat stress on the ovary

Ovaries are responsible for generating female gametes and producing hormones regulating reproductive functions [38]. All studies investigating the effects of increased temperature on the ovary were conducted *in vivo*, (n=9) (summarized in **Table 1**) in a range of species from mouse to cow. Temperatures tested ranged from 18°C – 33.3°C for the control and 31°C – 43°C for the HS groups. Exposure time to HS ranged from 30 minutes per day to upwards of 35 days. One group conducted measurements comparing shade vs. sun exposure [39].

**Table 1.**
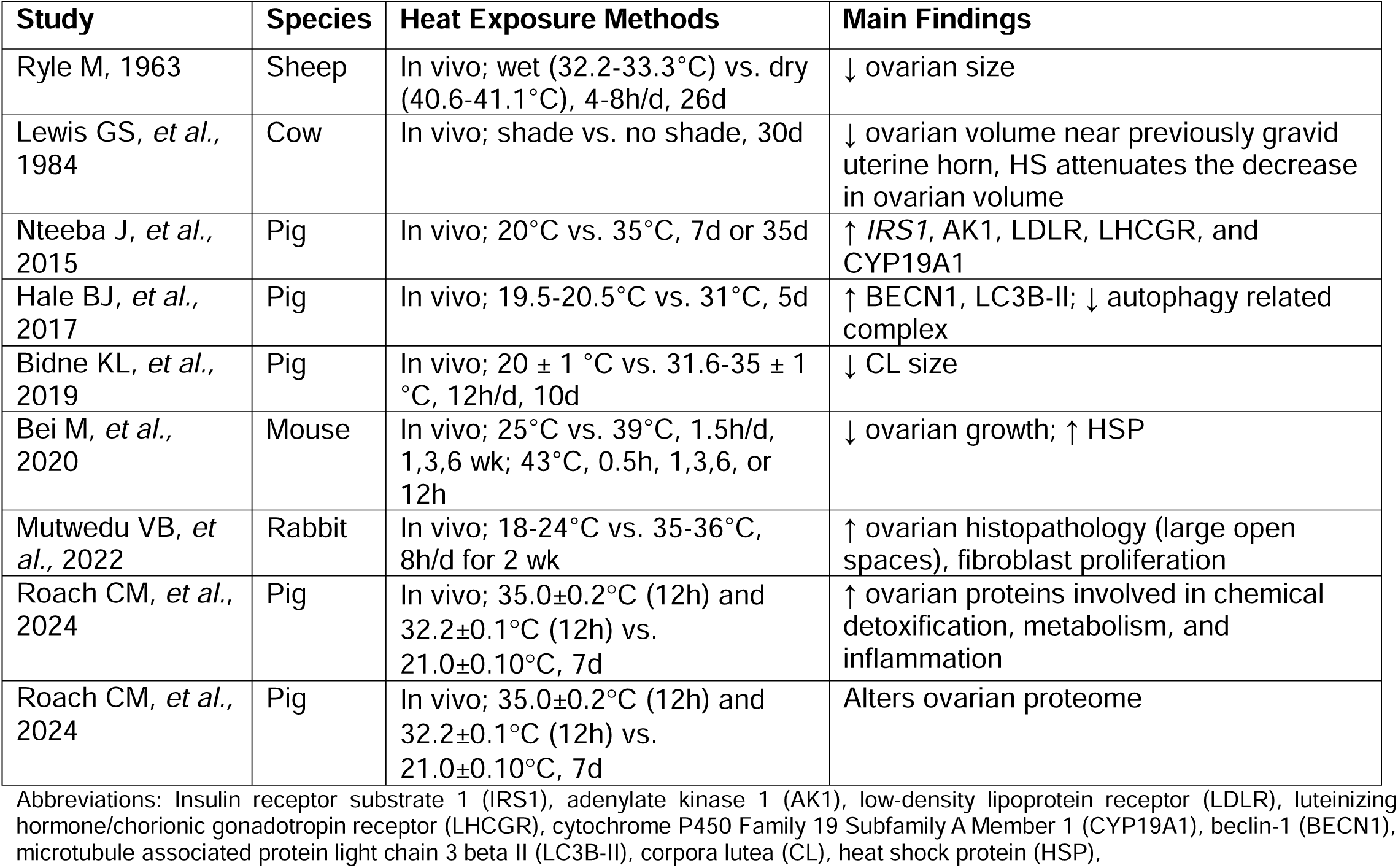
Effects of heat stress on the ovary.

Chronic HS in mice results in smaller ovaries with partially contracted central portion and reduced numbers of blood vessels, when animals are exposed to a daily 1.5-h session of 39°C at 50% humidity for durations of 1, 3, or 6 weeks, compared to controls kept at a constant temperature of 25°C with 50% humidity for the same durations [40]. In sheep, HS also reduces ovarian weight, with estimates of mean ovarian weight being 1344 mg in control compared to 1159 mg in the hot-room exposed animals [41]. Here, sheep were exposed to HS of 40.6°C for 4h/day starting on day 1 that was gradually increased to 41.1°C for 8h/day by day 9, compared to controls that were consistently exposed to 32.2 – 33.3°C. The study investigating the effects of shade compared to sun exposure assigned cows and heifers at 160-190 days of gestation to shade (n=7 cows; n=2 heifers) or no shade (n=8 cows; n=2 heifers), followed by examination of the reproductive tract 7 days postpartum [39]. Interestingly, in these cows, ovarian volume is reduced when ovaries are adjacent to the uterine horn that contained a fetus, and HS by sun exposure prepartum attenuates this effect [39]. HS in rabbits causes histopathological changes in the ovary, including focal areas of fibroblast proliferation in the interstitial tissue and the presence of large open spaces [42]. At the protein level, HS leads to proteins that are altered in the ovaries of control and HS pigs [43, 44], and they were noted to include those involved with chemical detoxification, metabolism, and inflammation [43]. At a molecular level, HS increases ovarian mRNA abundance of insulin receptor (IRS1), protein kinase B subunit 1 (AKT1), low-density lipoprotein receptor (LDLR), luteinizing hormone receptor (LHCGR), and aromatase (CYP19a) after 7 days exposure of gilts to 35°C compared to constant thermoneutral temperature of 20°C [45]. These results demonstrate that HS may alter ovarian insulin-mediated PI3K signaling, which plays a role in follicle activation and viability [45]. Additionally, HS in pigs induces autophagy in the ovary, marked by the upregulation of autophagy-related proteins including BECN1 and LC3-II, which are involved in the degradation of dysfunctional intracellular machinery [46]. Finally, corpora lutea (CLs) sizes in pigs are significantly reduced in the HS group compared to the controls [47]. Overall, these data indicate that HS leads to the reduction of ovarian size and weight, alters its histological architecture, and may change the expression of genes involved in autophagy and follicle function.

### The effects of heat stress on reproductive hormone levels

Studies investigating the effects of increased temperature on hormone levels in mammals were conducted *in vivo* (n=8), *in vitro* (n=4), and both *in vivo & in vitro* (n=2) (summarized in **Table 2**). Temperatures tested ranged from 15°C – 38.5°C for the control and 28°C – 43°C for the HS groups. Exposure time to HS ranged from 2h per day to upwards of 28 days. One group studied the retrospective analysis of meteorological data from 2003-2007 [48].

**Table 2.**
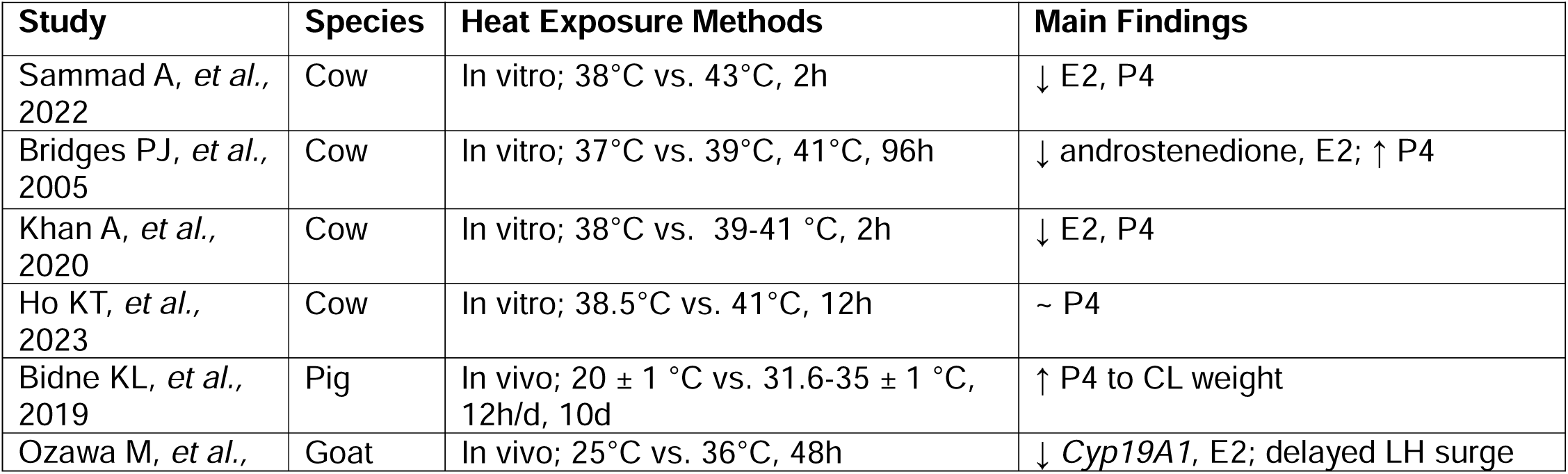
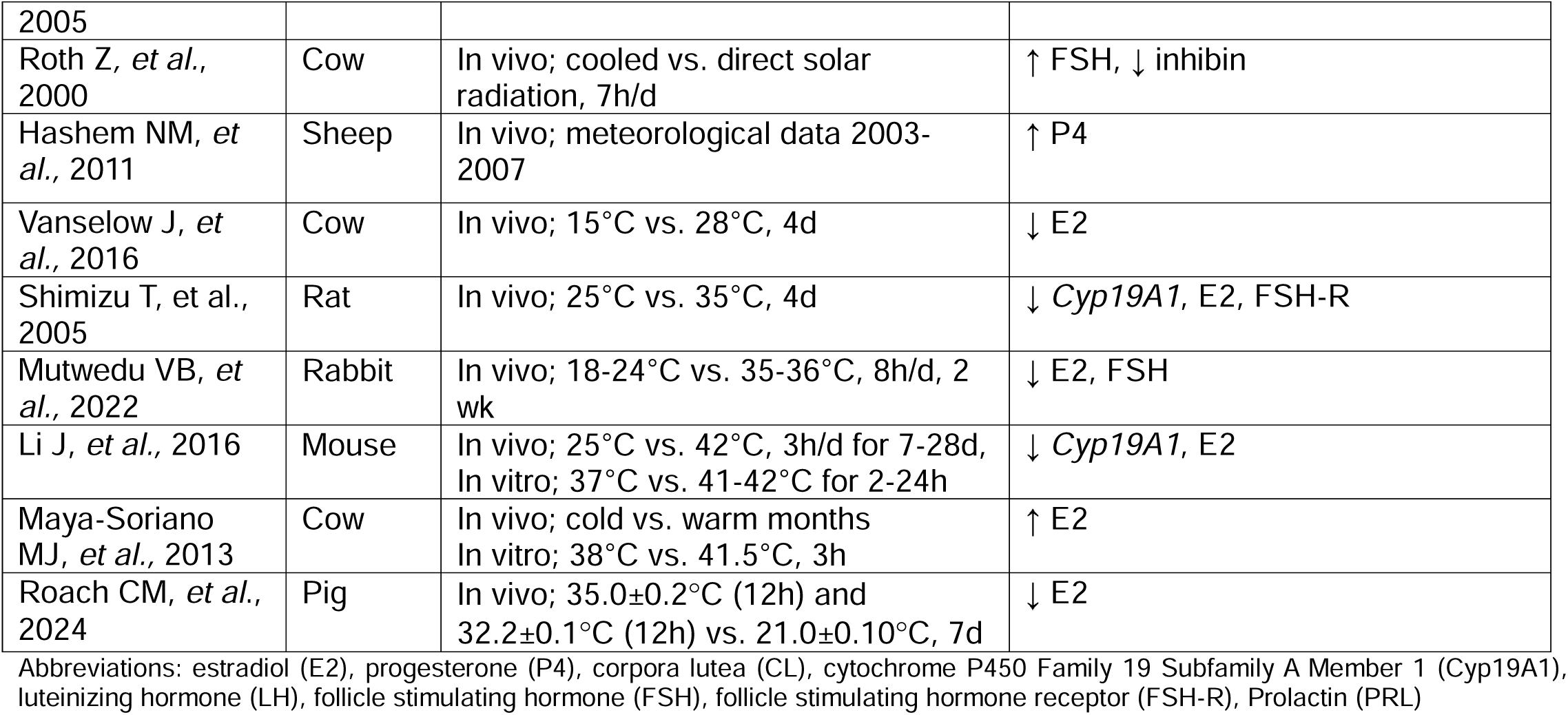
Effects of heat stress on reproductive hormones.

Some studies in a bovine model report that progesterone (P4) levels are not affected or are lower [49–51], while other studies in cows, pigs, and sheep find that P4 secretion is higher after HS exposure compared to controls [47, 48, 52]. Most studies demonstrate decreased estradiol (E2) levels in HS compared to control animals (mice, rats, rabbits, pigs, goats, and cows [42, 49, 50, 52–56]), with only one study in cows reporting an increase [57]. HS in cows also results in higher serum follicle stimulating hormone (FSH) levels and decreased inhibin concentrations during the heat exposed estrous cycle [58]. Additionally, HS decreases aromatase expression in granulosa cells in mice, rats, and goats [53–55]. Notably, HS in goats leads to a significant delay in the luteinizing hormone (LH) surge [54]. In addition, HS leads to increased levels of follistatin (FST), which may reduce FSH and LH levels, indirectly affecting the late growth of dominant follicles [59]; in rabbits, FSH levels decrease in response to HS [42]. Overall, these findings suggest that HS induces alterations in reproductive hormone levels. Most studies report a decrease in aromatase expression and E2 levels, with conflicting data on P4 and FSH. Further studies are warranted to examine whether the changes in ovarian hormone levels are due to direct effects of HS on the ovaries or secondary to the alterations in pituitary hormone secretion or both.

### The effects of heat stress on the follicles

Follicles represent the functional units of the ovary. They produce a mature oocyte and gonadal hormones contributing to the regulation of the ovulatory cycle [11]. Studies investigating the effects of increased temperature on the follicles were conducted *in vivo* (n=11), *in vitro* (n=3), and both *in vivo & in vitro* (n=2) (summarized in **Table 3**). Temperatures tested ranged from 15°C – 38.5°C for control and 28°C – 42°C for the HS groups. Exposure time to HS ranged from 2h to 30 days. Some groups performed seasonal measurements (i.e., winter vs summer collections), direct solar radiation vs. shade, or retrospectively examined ambient temperatures [39, 48, 58, 60–64].

**Table 3.**
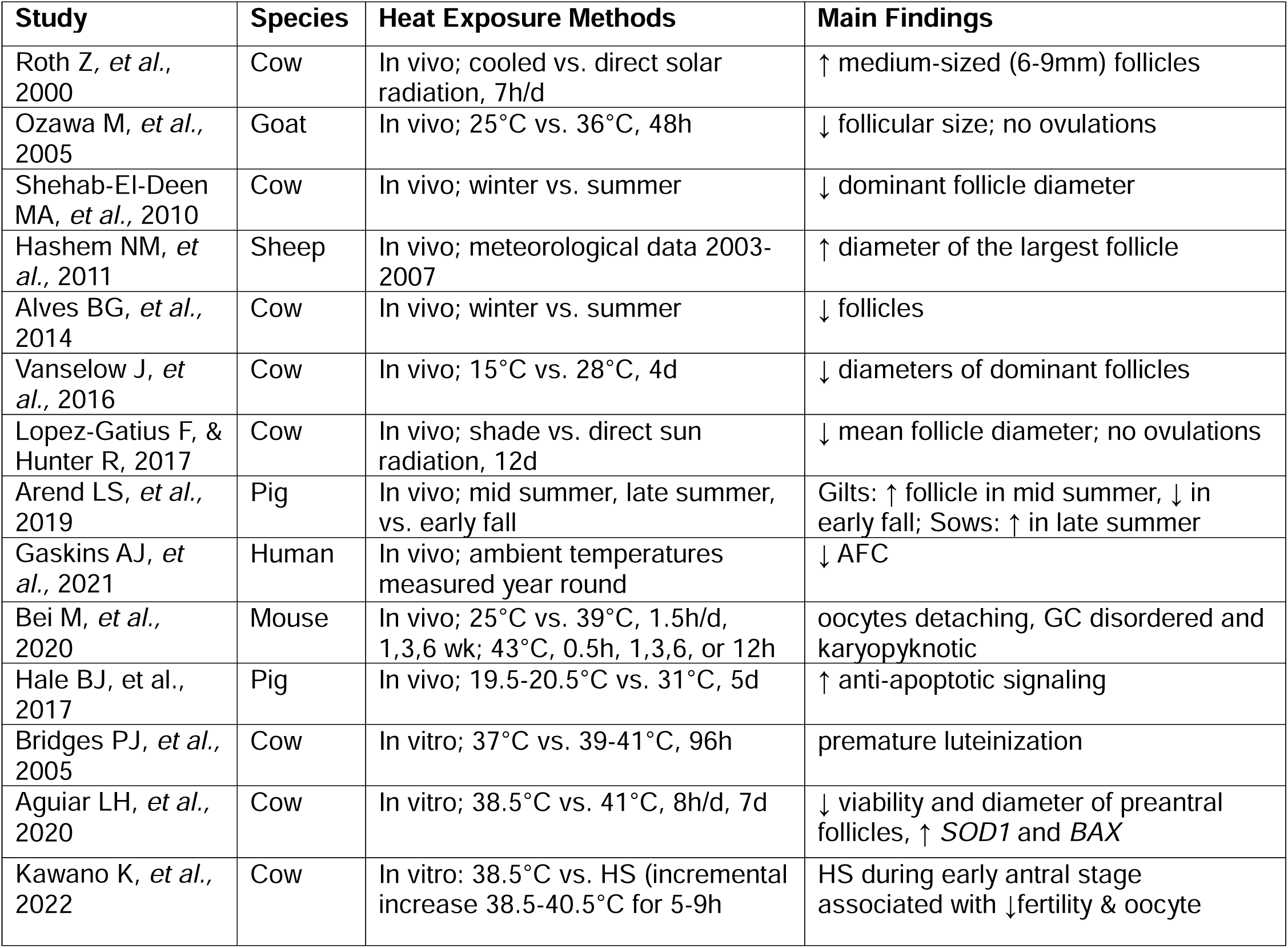
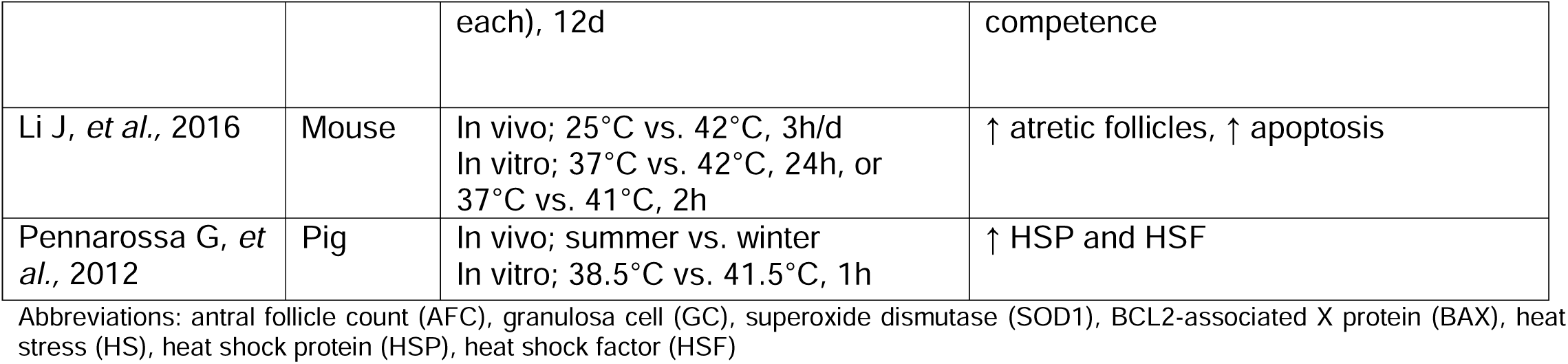
Effects of heat stress on follicles.

Interestingly, in cows, preovulatory follicles are cooler (0.74 ± 0.88°C) than neighboring uterine tissue and deep rectal temperatures under HS conditions suggesting a potential follicle-inherent thermoregulation mechanism [62]. HS in cows compromises folliculogenesis, with overall reduction in the size and number of follicles, fewer growing follicles, larger cohort of medium-sized follicles, and a decrease in the number of dominant follicles [58, 59, 61]. Preantral follicles, particularly those at the secondary stage (100.9 ± 13.7μm) of development, are the most sensitive to HS in cows, followed by early secondary (83.5 ± 8.4μm) and primary follicles (68.7 ± 9.2μm), indicating a stage-specific vulnerability to the increased temperatures [65]. Mechanistically, one study suggests that alterations of the activin-inhibin-follistatin system may partially be responsible for the tendency of compromised dominant follicle growth following HS [59].

Seasonal variations, perhaps at least partially mediated by temperature changes, influence the number of follicles in pigs. This results in the highest follicle numbers in mid-summer and the lowest in early fall [63]. In sheep, the diameter of the largest follicle is significantly higher during the breeding season (July) compared to winter (February) [48], which is in contrast to the dominant follicle diameter being smaller in summer compared to winter in cows [60]. Numerous studies in cows and goats also report a delayed deleterious effect of HS exposure on ovarian follicular growth [54, 58, 65]. In these models, HS reduces the viability and diameter of preantral follicles [54, 65]. In humans, a study in 631 subjects reports that higher ambient temperatures are linked to a lower antral follicle count (AFC), which is a measure of ovarian reserve. Specifically, a 1°C increase in average maximum temperature during the 90 days prior to ovarian reserve testing is associated with a 1.6% lower AFC. Additionally, this negative association is stronger November through June than during the summer months, suggesting the impact of possible acclimatization [64].

Evidence at histological and molecular level demonstrates that HS in mice leads to follicle atresia and dysregulated follicular development [40, 53], with oocytes detaching from the granulosa cell layer of antral follicles and granulosa cells becoming disordered and karyopyknotic [40]. Interestingly, HS in cows causes follicle cells to luteinize prematurely, which has been associated with persistent dominant follicles in cattle and accompanied by premature meiotic maturation of the oocyte, compromised subsequent embryo development, and fertility [52]. HS in pigs increases anti-apoptotic signaling in antral follicles [46]. Pig ovarian follicles constitutively express various heat shock proteins (HSPs) and heat shock factors (HSFs) in response to HS [66]. HSPs function to maintain cellular proteostasis and buffer against heat stressors [67]. HSFs are the transcription factors that regulate the expression of heat shock proteins [68]. In summary, folliculogenesis is significantly impaired in response to HS with most studies reporting a decrease in follicle numbers, size, and viability.

### The effects of heat stress on the oocyte

The oocyte is one of the largest cells in the body, is unique and extremely specialized, and undergoes dynamic processes during its maturation and development [69]. Most studies investigating the effects of increased temperature on the oocyte were conducted *in vitro* (n=38 *in vitro,* n=10 *in vivo*, n=5 *in vivo & in vitro*) (summarized in **Table 4**). From the 5 studies that examined HS *in vivo* and *in vitro,* 1 study looked at specific temperatures/duration *in vivo*, while the remaining 4 studies examined cold/winter vs. warm/summer months *in vivo*. Short term *in vivo* or *in vitro* exposure to HS (12h per day exposure to 32°C *in vivo* and 26h of maturation at 40°C *in vitro*) results in reduced porcine oocyte quality mediated in part by altered protein composition of the oocyte plasma membrane [70]. In cows, HS in summer months leads to impaired meiotic maturation, and compromised embryo cleavage, and blastocyst development rates [57, 71–73].

**Table 4.**
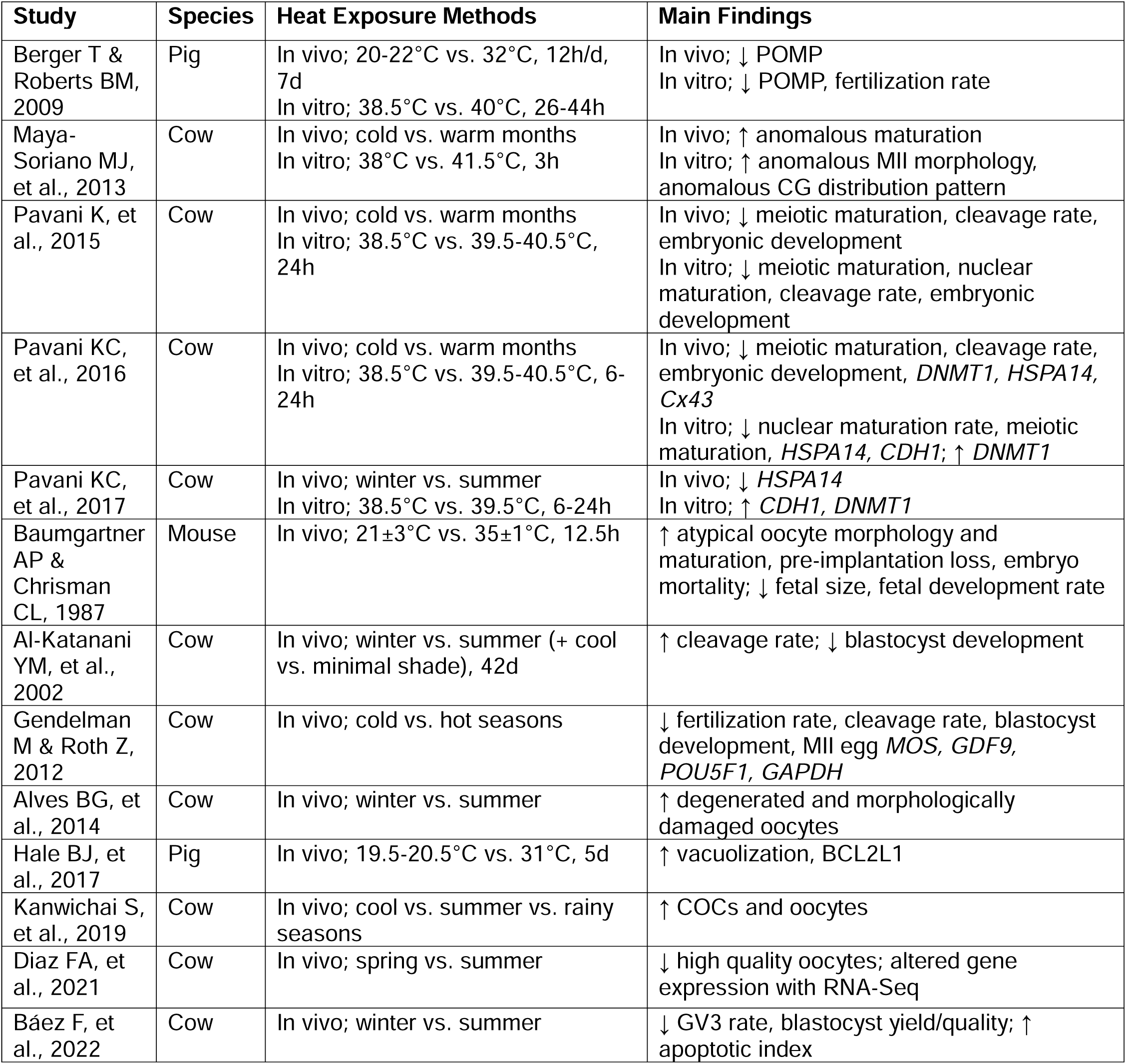
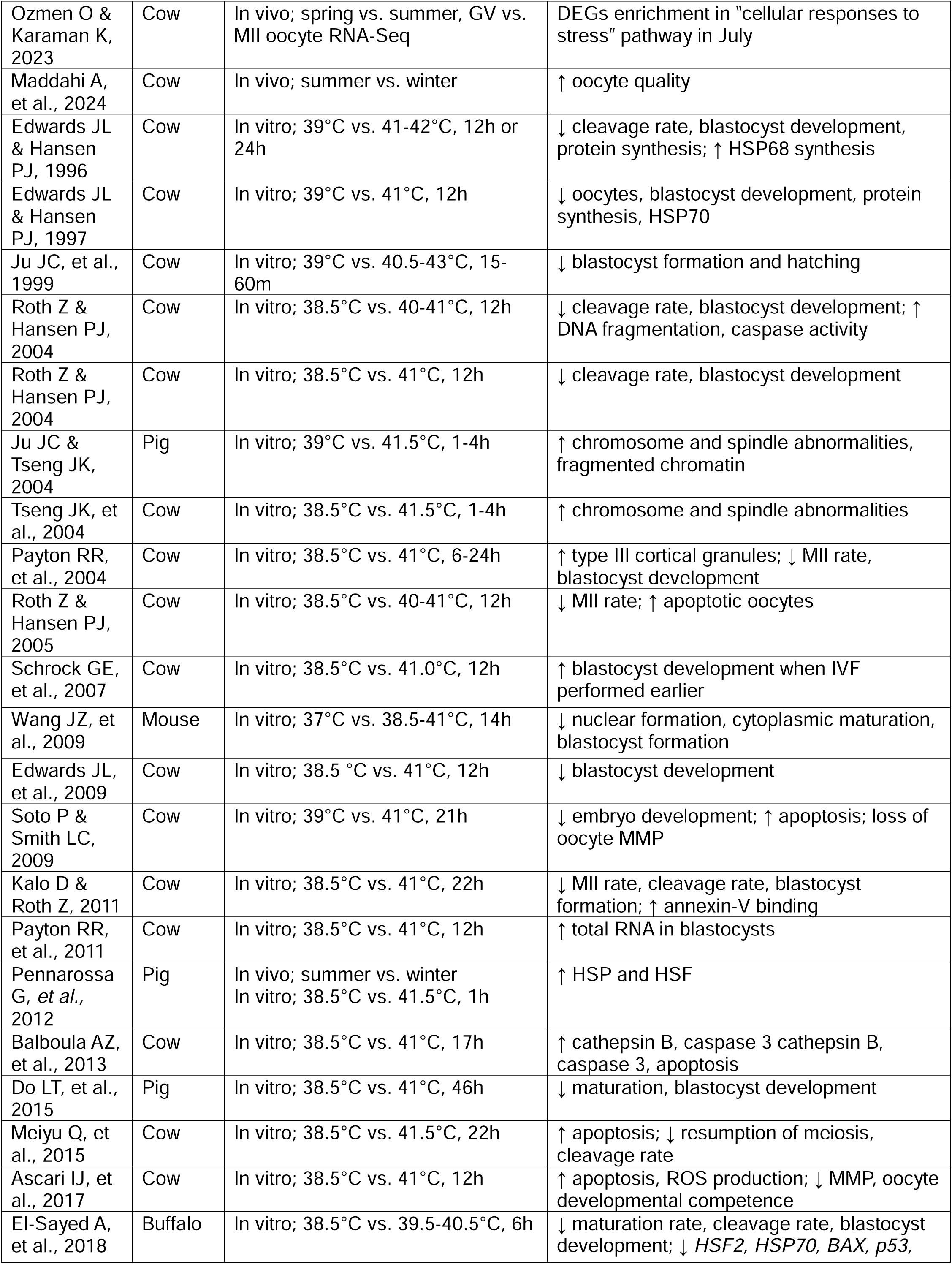
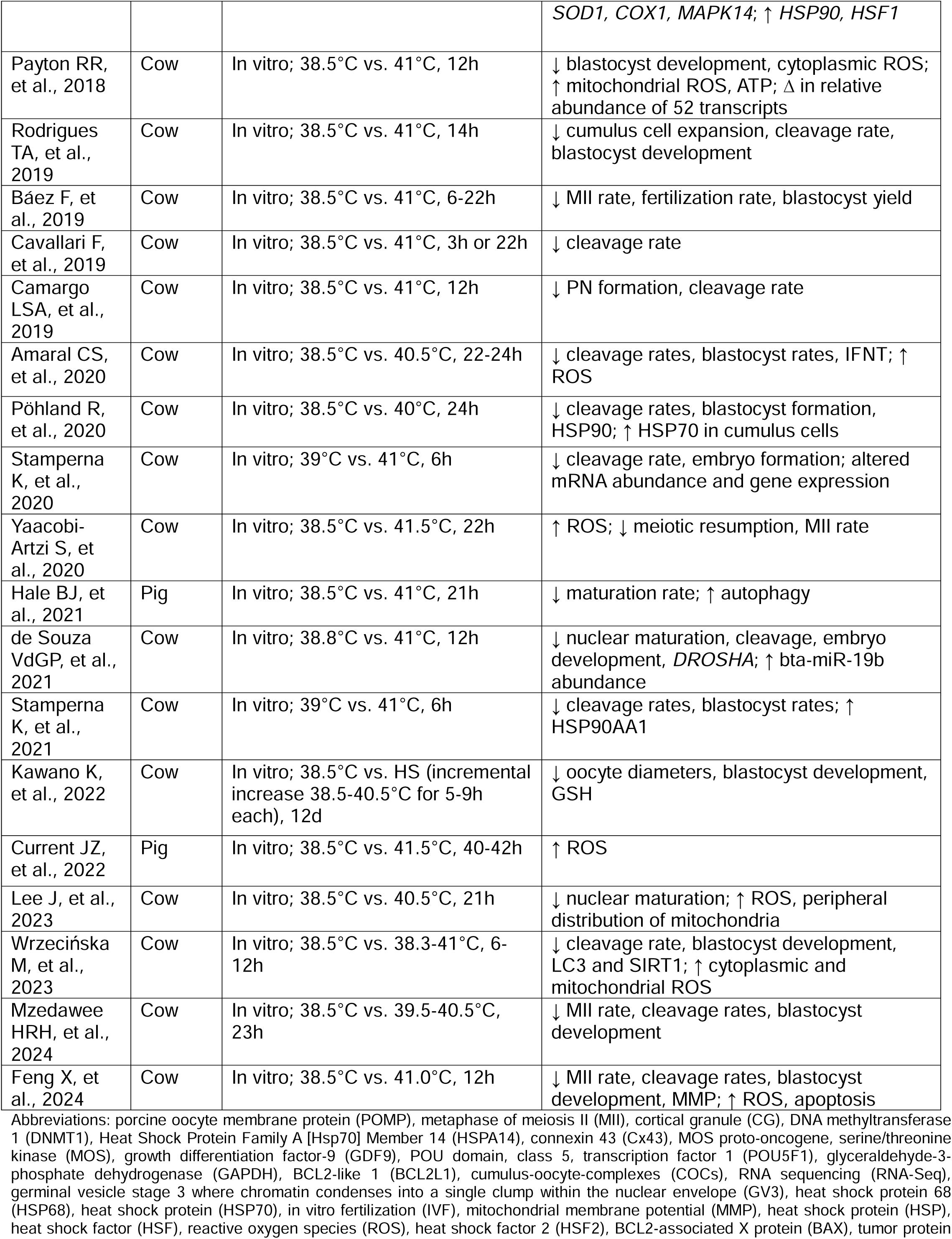
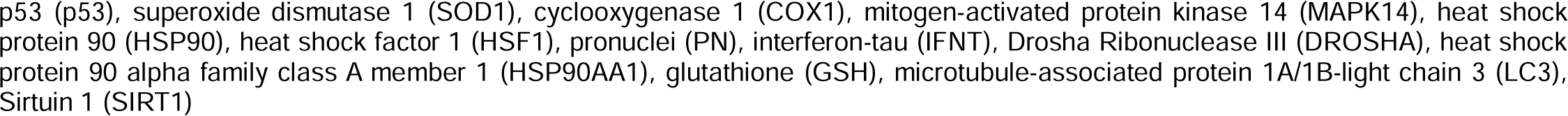
Effects of heat stress on the oocyte.

For the *in vivo* only studies, temperatures tested ranged from 19.5°C – 24°C for the control and 31°C – 36°C for the HS groups. Exposure time to HS ranged from 12.5h per day to 5 days. Most *in vivo* studies focus on seasonal temperature effects (n=7 out of 9). Seasonal HS in mice and cows leads to atypical oocyte morphologies, disrupted meiotic maturation, increase in the number of degenerated oocytes, and decrease in oocyte quality [61, 74, 75]. Seasonal HS in cows also leads to lower fertilization, embryo cleavage, and blastocyst development rates [76–78], with one study reporting that after HS exposure there is an increase in oocyte quality [79]. Results are conflicting in regard to HS impact on oocyte yield and *in vitro* maturation (IVM) in cows. Higher number of oocytes were obtained during the spring compared to summer in one study [75], whereas another study found collection numbers to be the highest during the summer [80]. After IVM, the percentage of mature, metaphase II (MII) eggs was not significantly different between seasons in one study [80]; however, another study found reduced oocyte maturation in the summer [77]. Lastly, HS in mice and pigs results in increased vacuolization in the oocyte, BCL21 protein abundance in the ovary, and decreased fetal size/development rate [46, 74]. Similar to the follicles, oocytes subjected to HS also exhibit delayed deleterious effects on gamete quality [78]. Embryo cleavage rates are decreased after *in vitro* fertilization (IVF) in cows [71]. At a molecular level, oocytes demonstrate significant changes in HSP and HSF expression levels in response to seasonal temperature variations [66]. In cow oocytes, when differentially expressed genes (DEGs) between germinal vesicle (GV) and MII stage oocytes were analyzed in May and July, only the July DEGs were enriched in “cellular responses to stress” pathway [81]. HS also increases anti-apoptotic signaling in pig oocytes [46].

For the *in vitro* studies, temperatures tested ranged from 37°C – 39°C for control groups and 38.5°C – 43°C for the HS groups. Exposure time to HS ranged from 15 minutes to 12 days. In cows, oocyte yield is decreased in HS compared control groups [82]. Additionally, oocyte-cumulus-granulosa complexes of cows exposed to HS (38.5-40.5°C) have significantly smaller mean oocyte diameters after IVM compared to the control group, indicating impaired oocyte growth due to heat exposure [83]. In a mouse study examining a range of elevated temperatures, oocyte maturation rates did not differ when exposed to 37-40°C. However, cytoplasmic maturation is impaired at 40°C, and nuclear formation is adversely affected beginning at 40.7°C [84]. Several studies in pig, cow, and buffalo did demonstrate decreased oocyte maturation with HS [85–94]. Similar to the *in vivo* studies, HS increases chromosome and spindle abnormalities in pig and cow oocytes *in vitro* [95, 96]. Three studies found that HS leads to the reduced oocyte mitochondrial membrane potential (MMP) [94, 97, 98]. Numerous studies report the decrease in embryo cleavage and blastocyst formation rates as a result of HS exposure in mice, pigs, and cows [82–84, 86, 89, 90, 93, 94, 98–113]. Interestingly, severe HS in cow oocytes at 43°C for up to 60 minutes during IVM followed by IVF showed that while blastocyst formation rates are not affected by HS for up to 30 minutes, longer exposures (45 and 60 minutes) significantly reduces both blastocyst and expanded blastocyst formation rates [104]. This indicates a threshold beyond which heat shock begins to negatively affect oocyte viability and developmental potential. Interestingly, cow blastocyst development rates improve when IVF is performed earlier in heat stressed eggs [114]. At a cellular and molecular level, cumulus-oocyte-complex (COC) HS exposure during IVM doesn’t change RNA abundance in oocytes, surrounding cumulus cells, or in 4- to 8- cell embryos but increases total RNA in resultant blastocysts after IVF [115]. Similar to the *in vivo* studies, several studies in pigs and cows demonstrate that HS during IVM induces apoptotic events in oocytes, as well as increases ROS and oxidative stress [91, 92, 97–99, 105, 106, 113, 116–118], with one study identifying members of the Bcl-2 family of peptides as having a protective effect [98]. Additionally, a possible breed-specific resilience to HS was reported. Compared to Jersey cows, HS in Holstein cows appears to adversely affect nuclear maturation at a higher degree, results in increased ROS levels, and abnormal oocyte mitochondria distribution suggesting impaired cytoplasmic maturation [118]. HS in cow oocytes at 41-41.5°C results in a significant increase in TUNEL-positive cells [97, 105, 116]. Antioxidant gene expression analysis in heat-stressed cow oocytes reveals that the expression GPX1, MnSOD, and G6PD tends to be upregulated [112, 119]. Overall, the evidence is overwhelming across species that HS detrimentally affects oocyte meiotic maturation, quality, and developmental competence both *in vivo* and *in vitro*.

### Effects of heat stress on the granulosa cells

Granulosa cells (GCs) support oocyte growth and development, ensure ovulation of developmentally competent gamete, and secrete hormones that regulate multiple organismal functions [120]. Studies that examined the impact of heat stress on GCs were conducted both *in vivo* and *in vitro* studies (n=4 *in vivo*, n=4 *in vitro,* n=1 *in vivo & in vitro*) (summarized in **Table 5**). For the *in vivo* studies, temperatures tested ranged from 15°C – 25°C for the control and 28°C – 42°C for the HS groups. Exposure time to HS ranged from 3h per day to 4 days. For the *in vitro* studies, temperatures ranged from 37°C – 38°C for the control and 39°C – 43°C for the HS groups. Exposure time to HS ranged from 2h to 24h.

**Table 5.** Effects of heat stress on granulosa cells.

| Study | Species | Heat Exposure Methods | Main Findings |
| --- | --- | --- | --- |
| Li J, et al., 2016 | Mouse | In vivo; 25°C vs. 42°C 3h/d<br>In vitro: 37°C vs. 42°C, 24h or 37°C vs. 41°C, 2h | ↑ apoptotic GCs; Caspase-3, Bim |
| Shimizu T, et al., 2005 | Rat | In vivo; 25°C vs. 35°C, 4d | ↑ apoptosis; ↑ *Bax* |
| Vanselow J, et al., 2016 | Cow | In vivo; 15°C vs. 28°C, 4d | ↑ *HAS3*, *CRHBP* |
| Hale BJ, et al., 2017 | Pig | In vivo; 19.5-20.5°C vs. 31°C, 5d | ↑ *BCL2L1*, vacuolization |
| Klabnik JL, et al., 2022 | Cow | In vivo; temperature humidity index (THI) of 65.8±0.1 vs. 71.0-86.4, 12.1±0.2h | 87 DEGs; DEGs suggested to promote ovulation, impact collagen formation or angiogenesis |
| Li L, et al., 2016 | Cow | In vitro; 38°C vs. 40.5°C, 2h | 1211 DEGs; ↑ apoptosis through BAX/BCL-2 pathway; ↓ steroidogenic gene mRNA expression |
| Khan A, et al., 2020 | Cow | In vitro; 38°C vs. 39°C, 40°C, 41°C, 2h | 142 DEGs (39°C); 321 DEGs (40°C); 294 DEGs (41°C); ↓ viability; ↑ apoptosis, ROS |
| Faheem MS, et al., 2021 | Buffalo | In vitro; 37°C vs. 39.5°C, 40.5°C, 41.5°C, 24h | ↓ viability, miR-1246, miR-181a, miR27b; ↑ *SOD2*, miR-708 |
| Sammad A, et al., 2022 | Cow | In vitro; 38°C vs. 43°C, 2h | 12,385 DEGs; ↑ ROS, apoptosis, stress-response genes |

Abbreviations: granulosa cells (GCs), B-cell Lymphoma 2-like protein 11 (Bim), BCL2-associated X protein (Bax), Hyaluronan Synthase 3 (*HAS3*), Corticotropin Releasing Hormone Binding Protein (*CRHBP*), BCL2-like 1 (*BCL2L1*), differentially expressed genes (DEGs), B-cell leukemia/lymphoma 2 (*BCL-2*), reactive oxygen species (ROS), superoxide dismutase 2 (*SOD2*)

HS in cows and buffalos reduces the viability of GCs compared to controls [50, 121]. HS in pigs increases the incidence of vacuolization in GCs [46]. HS exposure in mice and cows results in increased granulosa cell apoptosis [49, 50, 53, 122]. However, one study reported that HS in cows does not induce apoptosis in GCs but affects the expression of genes encoding the extracellular matrix proteins, glycoproteins, proteins playing a role in lipid synthesis, metabolic processes, and apoptosis [59]. HS in rats inhibits FSH-R expression in GCs, and this is associated with enhanced susceptibility to apoptosis [55]. At a molecular level, somatic components of the periovulatory follicle (i.e., cumulus cells surrounding the maturing oocyte and the mural GCs lining the follicle wall) demonstrate altered gene expression in response to varying degrees of elevated body temperatures, with majority of gene ontologies related to protein processing, the endoplasmic reticulum or Golgi apparatus [123]. Cow GCs cultured under HS (40.5°C) demonstrate altered gene expression with upregulated genes (*BCL-2, BAX, HSP*) involved in apoptosis regulation, and downregulated genes (*SF-1*, *CYP19A1, STAR, CYP11A1)* involved in secretion of E2 and P4 [122]. To summarize, HS in GCs reduces cell viability, increases apoptosis, and dysregulates gene expression.

### Interventions to mitigate the adverse effects of heat stress on granulosa cells, oocytes, and embryos

Studies investigating the effects of supplements to mitigate the sequelae of heat stress on mammalian GCs, oocytes, and embryos were conducted *in vivo* (n=7) and *in vitro* (n=15) (summarized in **Table 6**). Administration of Moringa oleifera aqueous seed extracts during HS in rabbits increases E2 and FSH [42]. Injection of pregnant mare serum gonadotropin (PMSG) during HS in rats results in decreased aromatase, E2, and FSH-R of GCs [55]. The addition of rapamycin during oocyte IVM in pigs under HS results in increased oocyte maturation rates [88]. Supplementation of FSH during IVM under HS in cows, improves oocyte maturation rates and embryo development [124]. Supplementation with astaxanthin during HS in pigs increases oocyte maturation, fertilization, and blastocyst development [86]. Addition of 10% follicular fluid to the oocyte maturation medium in cows, during IVM under HS, results in increased blastocyst development rates [108]. Treatment with Selenium during cow COC IVM under HS improves oocyte maturation and embryo cleavage rates, and the total number of blastocysts, as well as upregulates the expression of antioxidant genes (*CCND1, SEPP1, GPX-4, SOD, CAT*) [125]. Idebenone (IDB), potent antioxidant that functions as a carrier in electron transport chain, protects pig oocytes from HS during IVM (20-24 h, 42°C) through improving cumulus cell expansion, nuclear maturation, and blastocyst development rates [126]. Treatment with cysteine in COCs during IVM in cows under HS results in increased GSH levels and blastocyst development rates [83]. Addition of IGF-1 during IVM under HS in cows results in improved maturation, embryo cleavage rates, MMP, and decreases apoptotic cells [97, 105]. Treatment with anti-apoptotic peptides, BH4 domain of Bcl-xL (TAT-BH4), Bax inhibitor peptide (BIP), or combination of the two (BIP-BH4) decreases apoptosis and improves blastocyst development of heat stressed cow oocytes [98]. In cows, the addition of z-DEVD-fmk (group II caspase inhibitor) and sphingosine 1-phosphate blocks the negative effects of HS on embryo cleavage and subsequent development [91, 109, 110]. Supplementation of melatonin during HS in pigs increases total follicle numbers [63]. Supplementation with linoleic and linolenic acids significantly decreases ROS levels during pig oocyte IVM under HS conditions (40-42h under HS of 41.5°C) [117]. Additionally, in cows, treatment of GCs with a standardized extract of asparagus officinalis stem improves mitochondrial activity, reduces oxidative stress and increases the formation of lipid droplets, which are important for cholesterol storage used in steroid hormone production [51]. In pigs, while serum prolactin (PRL) did not differ from controls to HS only treated, when HS is supplemented with the addition of zearalenone (ZEN), there is significant increase in PRL [43]. In the same study, E2 was initially decreased with HS; however, when HS is supplemented with ZEN, there was a reduction in decrease of E2 [43]. In pigs, treatment of heat stressed COCs with Mogroside V attenuates deleterious effects of HS on oocyte maturation and blastocyst formation [127]. Additionally, Vitamin C supplementation to culture media in HS cow granulosa cells results in decreased ROS and apoptosis, and increased P4 and E2 [128]. Overall, investigators are seeking strategies to improve the detrimental impact of HS on mammalian oocytes and embryos. Although some compounds showed promise, further investigations and understanding of their mechanisms of actions are needed.

**Table 6.** Interventions to mitigate heat stress impacts on ovarian function.

| Study | Species | Heat Exposure Methods + Intervention | Main Findings |
| --- | --- | --- | --- |
| Mutwedu VB, et al., 2022 | Rabbit | In vivo; 18-24°C vs. 35-36°C, 8h/d, 2wk + *Moringa oleifera* aqueous seed extract | ↑ E2 & FSH |
| Ho KT, et al., 2023 | Cow | In vitro; 38.5°C vs. 41°C, 12h + EAS | ↑ P4; ↑ STAR, 3β-HSD, mitochondrial activity, lipid droplet formation; ↓ ROS |
| Shimizu T, et al., 2005 | Rat | In vivo; 25°C vs. 35°C, 4d + PMSG | ↓ aromatase activity, E2, FSH-R |
| Arend LS, et al., 2019 | Pig | In vivo; mid summer (26.2°C), late summer (28.3°C), vs. early fall (27.4°C) + melatonin | ↑ follicles |
| Maddahi A, et al., 2024 | Cow | In vivo; summer vs. winter + vitamins E, C, and coenzyme Q10 | ↑ maturation, cleavage rates |
| Roth Z & Hansen PJ, 2004 | Cow | In vitro; 38.5°C vs. 40- 41°C, 12h + z-DEVD-fmk | blocks HS effect on embryo cleavage rate and blastocyst development |
| Roth Z & Hansen PJ, 2004 | Cow | In vitro; 38.5°C vs. 41°C, 12h + S1P | blocks HS effect on cleavage and subsequent development |
| Roth Z & Hansen PJ, 2005 | Cow | In vitro; 38.5°C vs. 40- 41°C, 12h | S1P blocks the effect of HS on progression through meiosis |
| Soto P & Smith LC, 2009 | Cow | In vitro; 39°C vs. 41°C, 21h + BIP, TAT-BH4, or BIP+BH4 | ↑ HS oocytes development to blastocysts; ↓ embryonic development, ↓ apoptosis |
| Do LT, et al., 2015 | Pig | In vitro; 38.5°C vs. 41°C, 46h + astaxanthin | ↑ oocyte maturation, fertilization, blastocyst development; ↓ proportion of apoptotic cells in oocytes exposed to oxidative stress |
| Meiyu Q, et al., 2015 | Cow | In vitro; 38.5°C vs. 41.5°C, 22h + IGF-1 | ↑ proportion of MII-stage eggs and 4-cell-stage cleaved embryos |
| Ascari IJ, et al., 2017 | Cow | In vitro; 38.5°C vs. 41°C, 12h + IGF-1 | ↑ MMP, ROS production; ↓ apoptotic cells |
| Rodrigue s TA, et al., 2019 | Cow | In vitro; 38.5°C vs. 41°C, 14h + 10% follicular fluid | ↑ blastocyst development |
| Karaşahin T, 2020 | Cow | In vivo; autumn vs. spring + FSH | ↑ maturation & embryonic growth rates |
| Hale BJ, et al., 2021 | Pig | In vitro; 38.5°C vs. 41°C, 21h + rapamycin | ↑ the maturation rate |
| Shi XY, et al., 2022 | Pig | In vitro; 38.5°C vs. 42°C, 4h + IDB | ↑ quality and developmental competence of oocytes, antioxidant capacity, GDF9 and BMP15; protects mitochondrial function |
| Kawano K, et al., 2022 | Cow | In vitro; 38.5°C control vs HS (incremental increase 38.5-40.5°C for 5-9h each), 12d + cysteine | ↑ GSH levels, blastocyst rate |
| Current JZ, et al., 2022 | Pig | In vitro; 38.5°C vs. 41.5°C, 40-42h + linoleic and linolenic acids | ↓ ROS; ↑ fertilization rates, cleavage rates, blastocyst formation rates, antioxidative enzymes (SOD and catalase), GSH |
| Toosinia S, et al., 2023 | Cow | In vitro; 38.5°C vs. 41°C HS, 24h + Se | ↑ viability of cumulus cells and oocytes, MII rate, cleavage rate, blastocyst development, antioxidative defense genes; ↓ apoptosis-related genes |
| Roach CM, et al., 2024 | Pig | In vivo; 35.0±0.2°C (12h) and 32.2±0.1°C (12h) vs. 21.0±0.10°C, 7d + ZEN | ↑ Prolactin, ↑E2 |
| Peng K, et al., 2024 | Pig | In vitro; 38.5°C vs. 42°C + MV | ↑ MII rate, blastocyst rate |
| Sammad A, et al., 2024 | Cow | In vitro; 38°C vs. 43°C, 2h + Vitamin C | ↓ ROS, apoptosis; ↑ P4, E2 |

Abbreviations: Estradiol (E2), follicle stimulating hormone (FSH), Extract of Asparagus officinalis stem (EAS), progesterone (P4), steroidogenic acute regulatory protein (STAR), 3 beta-hydroxysteroid dehydrogenase (3β-HSD), reactive oxygen species (ROS), pregnant mare serum gonadotropin (PMSG), follicle stimulating hormone receptor (FSH-R), caspase-3 inhibitor (z-DEVD-fmk), heat stress (HS), sphingosine 1-phosphate (S1P), anti-apoptotic peptides BH4 domain of Bcl-xL (TAT-BH4), Bax inhibitor peptide (BIP), insulin-like growth factor 1 (IGF-1), metaphase of meiosis II (MII), mitochondrial membrane potential (MMP), idebenone (IDB), glutathione (GSH), selenium (Se), zearalenone (ZEN), Mogroside V (MV)

## Conclusions

There are significant adverse impacts of HS on the ovaries, reproductive hormone profiles, follicles, oocytes, and GCs in mammals (**Figure 2**). HS results in dysregulated gene expression and increases apoptosis in multiple ovarian cell types. In the ovary, HS reduces ovarian size and weight, disrupts histology, and induces autophagy. In the follicles, HS leads to the increased follicle atresia, reduced size and number of follicles, and a decrease in the number of dominant follicles. In the oocytes, HS negatively impacts oocyte growth, maturation, and developmental competence. In the GCs, HS results in significantly decreased viability and increased apoptosis. These findings underscore the multifaceted impact of HS on ovarian function. With the projected increase in high-temperature days in the coming decades, elucidating the mechanisms by which HS affects the ovary is critical. Current knowledge of the molecular pathways involved remains limited, necessitating further research to inform the development of effective therapeutic interventions. Efforts to identify compounds that mitigate HS-induced damage to oocytes and embryos are ongoing, highlighting the urgency of advancing strategies to preserve reproductive health under thermal stress.

**Figure 2.**
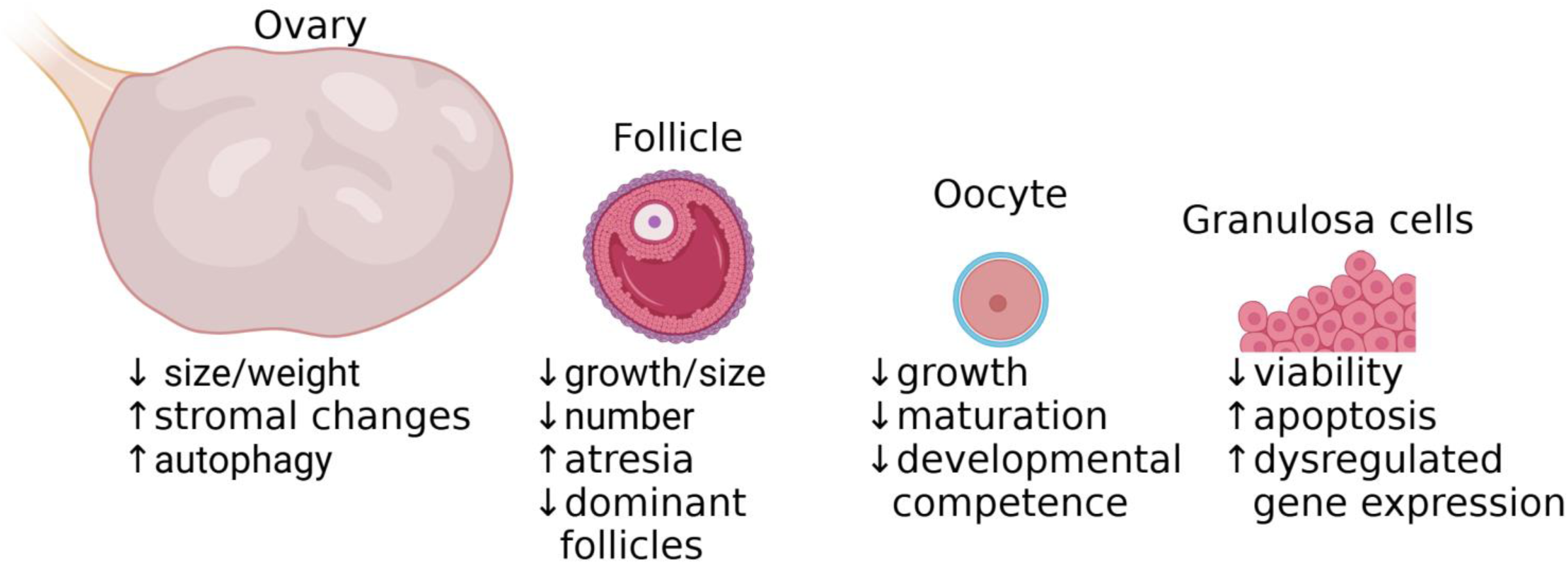
Schematic summarizing the evidence on heat stress effects on the ovary, follicle, oocyte, and granulosa cells.

## Notes

**Grant Support:** This project was supported by Women’s Reproductive Health Research (WRHR) Career Development Program K12 HD050121 (EB)

### Competing Interest Statement

The authors have declared no competing interest.

## References

1. Romanello, M., et al., The 2021 report of the Lancet Countdown on health and climate change: code red for a healthy future. Lancet, 2021. 398(10311): p. 1619-1662.

2. IPCC Sixth Assessment Report. 2022.

3. Climate Effects on Health. 2022; Available from: https://www.cdc.gov/climateandhealth/effects/default.htm.

4. Climate Change. Available from: https://www.who.int/health-topics/climate-change#tab=tab_1.

5. Walton, D.v.A., M., Climate-related extreme weather events and COVID-19. 2020.

6. Segal, T.R. and L.C. Giudice, Systematic review of climate change effects on reproductive health. Fertil Steril, 2022. 118(2): p. 215–223.

7. Perez-Crespo, M., B. Pintado, and A. Gutierrez-Adan, Scrotal heat stress effects on sperm viability, sperm DNA integrity, and the offspring sex ratio in mice. Mol Reprod Dev, 2008. 75(1): p. 40–7.

8. Meyerhoeffer, D.C., et al., Reproductive criteria of beef bulls during and after exposure to increased ambient temperature. J Anim Sci, 1985. 60(2): p. 352–7.

9. Houston, B.J., et al., Heat exposure induces oxidative stress and DNA damage in the male germ line. Biol Reprod, 2018. 98(4): p. 593–606.

10. Macklon, N.S. and B.C. Fauser, Follicle development during the normal menstrual cycle. Maturitas, 1998. 30(2): p. 181–8.

11. McGee, E.A. and A.J. Hsueh, Initial and cyclic recruitment of ovarian follicles. Endocr Rev, 2000. 21(2): p. 200–14.

12. Duncan, F.E. and J.L. Gerton, Mammalian oogenesis and female reproductive aging. Aging (Albany NY), 2018. 10(2): p. 162–163.

13. Minkin, M.J., *Menopause: Hormones, Lifestyle,* and Optimizing Aging. Obstet Gynecol Clin North Am, 2019. 46(3): p. 501–514.

14. van Dijk, G.M., et al., Health issues for menopausal women: the top 11 conditions have common solutions. Maturitas, 2015. 80(1): p. 24–30.

15. Takahashi, T.A. and K.M. Johnson, Menopause. Med Clin North Am, 2015. 99(3): p. 521–34.

16. Magnus, M.C., et al., Role of maternal age and pregnancy history in risk of miscarriage: prospective register based study. BMJ, 2019. 364: p. l869.

17. American College of, O., P. Gynecologists Committee on Gynecologic, and C. Practice, *Female age-related fertility decline. Committee Opinion No. 589*. Fertil Steril, 2014. 101(3): p. 633-4.

18. Hansen, J., et al., Global temperature change. Proc Natl Acad Sci U S A, 2006. 103(39): p. 14288–93.

19. Davariashtiyani, A., et al., Exponential increases in high-temperature extremes in North America. Sci Rep, 2023. 13(1): p. 19177.

20. Ouzzani, M., et al., Rayyan—a web and mobile app for systematic reviews. Systematic Reviews, 2016. 5(1): p. 210.

21. Pollard, J.W., et al., Effect of ambient temperatures during oocyte recovery on in vitro production of bovine embryos. Theriogenology, 1996. 46(5): p. 849–58.

22. Nasiri, A.H., et al., Effects of live yeast dietary supplementation on hormonal profile, ovarian follicular dynamics, and reproductive performance in dairy cows exposed to high ambient temperature. Theriogenology, 2018. 122: p. 41–46.

23. El-Ratel, I.T., A.E. Abdel-Khalek, and S.F. Fouda, Effect of ovarian stimulation by different gonadotrophin treatments on in vivo and in vitro reproductive efficiency of rabbit does under high ambient temperature. Trop Anim Health Prod, 2020. 53(1): p. 22.

24. Khan, A., et al., SOD1 Gene Silencing Promotes Apoptosis and Suppresses Proliferation of Heat-Stressed Bovine Granulosa Cells via Induction of Oxidative Stress. Vet Sci, 2021. 8(12).

25. Rhouma, R. and M. Aroua, Super-ovulatory response of pregnant mare serum gonadotropin (PMSG) treatment on ovarian characteristics of rabbit does under heat stress. The Indian Journal of Animal Sciences, 2022. 92.

26. Roth, Z., et al., Delayed effect of heat stress on steroid production in medium-sized and preovulatory bovine follicles. Reproduction, 2001. 121(5): p. 745–51.

27. Roth, Z., et al., Carry-over effect of summer thermal stress on characteristics of the preovulatory follicle of lactating cows. Journal of Thermal Biology, 2004. 29(7): p. 681–685.

28. de, S.T.-J.J.R., et al., Effect of maternal heat-stress on follicular growth and oocyte competence in Bos indicus cattle. Theriogenology, 2008. 69(2): p. 155–66.

29. Graves, K.L., et al., Characterizing the acute heat stress response in gilts: II. Assessing repeatability and association with fertility. J Anim Sci, 2018. 96(6): p. 2419–2426.

30. Roth, Z., et al., Improvement of quality of oocytes collected in the autumn by enhanced removal of impaired follicles from previously heat-stressed cows. Reproduction, 2001. 122(5): p. 737–44.

31. Roth, Z., et al., Effect of treatment with follicle-stimulating hormone or bovine somatotropin on the quality of oocytes aspirated in the autumn from previously heat-stressed cows. J Dairy Sci, 2002. 85(6): p. 1398–405.

32. Vendrell-Flotats, M., et al., Effect of heat stress during in vitro maturation on developmental competence of vitrified bovine oocytes. Reprod Domest Anim, 2017. 52 **Suppl 4**: p. 48–51.

33. Khan, A., et al., RNAi-Mediated Silencing of Catalase Gene Promotes Apoptosis and Impairs Proliferation of Bovine Granulosa Cells under Heat Stress. Animals (Basel), 2020. 10(6).

34. Mishra, V., A.K. Misra, and R. Sharma, Effect of ambient temperature on in vitro fertilization of Bubaline oocyte. Anim Reprod Sci, 2007. 100(3-4): p. 379–84.

35. Wang, Y.R., et al., Heme oxygenase 1 regulates apoptosis induced by heat stress in bovine ovarian granulosa cells via the ERK1/2 pathway. J Cell Physiol, 2019. 234(4): p. 3961–3972.

36. Feltes, G.L., et al., Impact of heat stress on genetic evaluation of oocyte and embryo production in Gir dairy cattle. Trop Anim Health Prod, 2023. 56(1): p. 7.

37. Contreras-Mendez, L.A., et al., The Anti-Mullerian Hormone as Endocrine and Molecular Marker Associated with Reproductive Performance in Holstein Dairy Cows Exposed to Heat Stress. Animals (Basel), 2024. 14(2).

38. Puttabyatappa, M. and V. Padmanabhan, Chapter Fourteen - Developmental Programming of Ovarian Functions and Dysfunctions, in Vitamins and Hormones, G. Litwack, Editor. 2018, Academic Press. p. 377-422.

39. Lewis, G.S., et al., Effects of heat stress during pregnancy on postpartum reproductive changes in Holstein cows. J Anim Sci, 1984. 58(1): p. 174–86.

40. Bei, M., et al., Effects of heat stress on ovarian development and the expression of HSP genes in mice. J Therm Biol, 2020. 89: p. 102532.

41. Ryle, M., Early reproductive failure of ewes in a hot environment IV. The ovary. The Journal of Agricultural Science, 1963. 60(1): p. 101–104.

42. Mutwedu, V.B., et al., Effects of Moringa oleifera aqueous seed extracts on reproductive traits of heat-stressed New Zealand white female rabbits. Front Vet Sci, 2022. 9: p. 883976.

43. Roach, C.M., et al., Zearalenone exposure differentially affects the ovarian proteome in pre- pubertal gilts during thermal neutral and heat stress conditions. J Anim Sci, 2024. 102.

44. Roach, C.M., et al., Heat stress alters the ovarian proteome in prepubertal gilts. J Anim Sci, 2024. 102.

45. Nteeba, J., et al., Heat Stress Alters Ovarian Insulin-Mediated Phosphatidylinositol-3 Kinase and Steroidogenic Signaling in Gilt Ovaries. Biol Reprod, 2015. 92(6): p. 148.

46. Hale, B.J., et al., Heat stress induces autophagy in pig ovaries during follicular development. Biol Reprod, 2017. 97(3): p. 426–437.

47. Bidne, K.L., et al., Heat stress during the luteal phase decreases luteal size but does not affect circulating progesterone in gilts1. J Anim Sci, 2019. 97(10): p. 4314–4322.

48. Hashem, N.M., et al., Effect of season, month of parturition and lactation on estrus behavior and ovarian activity in Barki x Rahmani crossbred ewes under subtropical conditions. Theriogenology, 2011. 75(7): p. 1327–35.

49. Sammad, A., et al., Transcriptome Reveals Granulosa Cells Coping through Redox, Inflammatory and Metabolic Mechanisms under Acute Heat Stress. Cells, 2022. 11(9).

50. Khan, A., et al., Evaluation of heat stress effects on cellular and transcriptional adaptation of bovine granulosa cells. J Anim Sci Biotechnol, 2020. 11: p. 25.

51. Ho, K.T., et al., Synergistic effect of standardized extract of Asparagus officinalis stem and heat shock on progesterone synthesis with lipid droplets and mitochondrial function in bovine granulosa cells. J Steroid Biochem Mol Biol, 2023. 225: p. 106181.

52. Bridges, P.J., M.A. Brusie, and J.E. Fortune, Elevated temperature (heat stress) in vitro reduces androstenedione and estradiol and increases progesterone secretion by follicular cells from bovine dominant follicles. Domest Anim Endocrinol, 2005. 29(3): p. 508–22.

53. Li, J., et al., Effects of chronic heat stress on granulosa cell apoptosis and follicular atresia in mouse ovary. J Anim Sci Biotechnol, 2016. 7: p. 57.

54. Ozawa, M., et al., Alterations in follicular dynamics and steroidogenic abilities induced by heat stress during follicular recruitment in goats. Reproduction, 2005. 129(5): p. 621–30.

55. Shimizu, T., et al., Heat stress diminishes gonadotropin receptor expression and enhances susceptibility to apoptosis of rat granulosa cells. Reproduction, 2005. 129(4): p. 463–72.

56. Roach, C.M., et al., Phenotypic, endocrinological, and metabolic effects of zearalenone exposure and additive effect of heat stress in prepubertal female pigs. J Therm Biol, 2024. 119: p. 103742.

57. Maya-Soriano, M.J., et al., Bovine oocytes show a higher tolerance to heat shock in the warm compared with the cold season of the year. Theriogenology, 2013. 79(2): p. 299–305.

58. Roth, Z., et al., Immediate and delayed effects of heat stress on follicular development and its association with plasma FSH and inhibin concentration in cows. J Reprod Fertil, 2000. 120(1): p. 83–90.

59. Vanselow, J., et al., Exposure of Lactating Dairy Cows to Acute Pre-Ovulatory Heat Stress Affects Granulosa Cell-Specific Gene Expression Profiles in Dominant Follicles. PLoS One, 2016. 11(8): p. e0160600.

60. Shehab-El-Deen, M.A., et al., Biochemical changes in the follicular fluid of the dominant follicle of high producing dairy cows exposed to heat stress early post-partum. Anim Reprod Sci, 2010. 117(3-4): p. 189–200.

61. Alves, B.G., et al., Ovarian activity and oocyte quality associated with the biochemical profile of serum and follicular fluid from Girolando dairy cows postpartum. Anim Reprod Sci, 2014. 146(3-4): p. 117–25.

62. Lopez-Gatius, F. and R. Hunter, Clinical relevance of pre-ovulatory follicular temperature in heat-stressed lactating dairy cows. Reprod Domest Anim, 2017. 52(3): p. 366–370.

63. Arend, L.S., et al., Effects of feeding melatonin during proestrus and early gestation to gilts and parity 1 sows to minimize effects of seasonal infertility1. J Anim Sci, 2019. 97(11): p. 4635–4646.

64. Gaskins, A.J., et al., Impact of ambient temperature on ovarian reserve. Fertil Steril, 2021. 116(4): p. 1052–1060.

65. Aguiar, L.H., et al., Heat stress impairs in vitro development of preantral follicles of cattle. Anim Reprod Sci, 2020. 213: p. 106277.

66. Pennarossa, G., et al., Characterization of the constitutive pig ovary heat shock chaperone machinery and its response to acute thermal stress or to seasonal variations. Biol Reprod, 2012. 87(5): p. 119.

67. Hu, C., et al., Heat shock proteins: Biological functions, pathological roles, and therapeutic opportunities. MedComm (2020), 2022. 3(3): p. e161.

68. Akerfelt, M., R.I. Morimoto, and L. Sistonen, Heat shock factors: integrators of cell stress, development and lifespan. Nat Rev Mol Cell Biol, 2010. 11(8): p. 545–55.

69. Li, M., et al., Chapter 13 - Gene Expression During Oogenesis and Oocyte Development, in *The Ovary (Third Edition)*, P.C.K. Leung and E.Y. Adashi, Editors. 2019, Academic Press. p. 205-216.

70. Berger, T. and B.M. Roberts, Reduced immunolabelling of a porcine oocyte membrane protein reflects reduced fertilizability of porcine oocytes following elevated ambient temperature. Reprod Domest Anim, 2009. 44(2): p. 260–5.

71. Pavani, K., et al., Reproductive Performance of Holstein Dairy Cows Grazing in Dry-summer Subtropical Climatic Conditions: Effect of Heat Stress and Heat Shock on Meiotic Competence and In vitro Fertilization. Asian-Australas J Anim Sci, 2015. 28(3): p. 334–42.

72. Pavani, K.C., et al., Gene expression, oocyte nuclear maturation and developmental competence of bovine oocytes and embryos produced after in vivo and in vitro heat shock. Zygote, 2016. 24(5): p. 748–59.

73. Pavani, K.C., et al., The effect of kinetic heat shock on bovine oocyte maturation and subsequent gene expression of targeted genes. Zygote, 2017. 25(3): p. 383–389.

74. Baumgartner, A.P. and C.L. Chrisman, Embryonic mortality caused by maternal heat stress during mouse oocyte maturation. Animal Reproduction Science, 1987. 14(4): p. 309–316.

75. Diaz, F.A., et al., Evaluation of Seasonal Heat Stress on Transcriptomic Profiles and Global DNA Methylation of Bovine Oocytes. Front Genet, 2021. 12: p. 699920.

76. Al-Katanani, Y.M., F.F. Paula-Lopes, and P.J. Hansen, Effect of season and exposure to heat stress on oocyte competence in Holstein cows. J Dairy Sci, 2002. 85(2): p. 390–6.

77. Baez, F., et al., Effect of season on germinal vesicle stage, quality, and subsequent in vitro developmental competence in bovine cumulus-oocyte complexes. J Therm Biol, 2022. 103: p. 103171.

78. Gendelman, M. and Z. Roth, Seasonal effect on germinal vesicle-stage bovine oocytes is further expressed by alterations in transcript levels in the developing embryos associated with reduced developmental competence. Biol Reprod, 2012. 86(1): p. 1–9.

79. Maddahi, A., et al., Exploring the impact of heat stress on oocyte maturation and embryo development in dairy cattle using a culture medium supplemented with vitamins E, C, and coenzyme Q10. J Therm Biol, 2024. **119**: p. 103759.

80. Kanwichai, S., et al., In vitro maturation of class I oocytes of bovine during different tropical seasons. Trop Anim Health Prod, 2019. 51(5): p. 1279–1282.

81. Ozmen, O. and K. Karaman, Transcriptome analysis and potential mechanisms of bovine oocytes under seasonal heat stress. Anim Biotechnol, 2023. 34(4): p. 1179–1195.

82. Edwards, J.L. and P.J. Hansen, Differential responses of bovine oocytes and preimplantation embryos to heat shock. Mol Reprod Dev, 1997. 46(2): p. 138–45.

83. Kawano, K., et al., Effect of heat exposure on the growth and developmental competence of bovine oocytes derived from early antral follicles. Sci Rep, 2022. 12(1): p. 8857.

84. Wang, J.Z., et al., Effects of heat stress during in vitro maturation on cytoplasmic versus nuclear components of mouse oocytes. Reproduction, 2009. 137(2): p. 181–9.

85. Baez, F., et al., Time-dependent effects of heat shock on the zona pellucida ultrastructure and in vitro developmental competence of bovine oocytes. Reprod Biol, 2019. 19(2): p. 195–203.

86. Do, L.T., et al., Astaxanthin present in the maturation medium reduces negative effects of heat shock on the developmental competence of porcine oocytes. Reprod Biol, 2015. 15(2): p. 86–93.

87. El-Sayed, A., et al., Developmental and molecular responses of buffalo (Bubalus bubalis) cumulus-oocyte complex matured in vitro under heat shock conditions. Zygote, 2018. 26(2): p. 177–190.

88. Hale, B.J., et al., Characterization of the effects of heat stress on autophagy induction in the pig oocyte. Reprod Biol Endocrinol, 2021. 19(1): p. 107.

89. Kalo, D. and Z. Roth, Involvement of the sphingolipid ceramide in heat-shock-induced apoptosis of bovine oocytes. Reprod Fertil Dev, 2011. 23(7): p. 876–88.

90. Payton, R.R., et al., Susceptibility of bovine germinal vesicle-stage oocytes from antral follicles to direct effects of heat stress in vitro. Biol Reprod, 2004. 71(4): p. 1303–8.

91. Roth, Z. and P.J. Hansen, Disruption of nuclear maturation and rearrangement of cytoskeletal elements in bovine oocytes exposed to heat shock during maturation. Reproduction, 2005. 129(2): p. 235–44.

92. Yaacobi-Artzi, S., et al., Melatonin slightly alleviates the effect of heat shock on bovine oocytes and resulting blastocysts. Theriogenology, 2020. 158: p. 477–489.

93. Mzedawee, H.R.H., et al., Heat shock interferes with the amino acid metabolism of bovine cumulus-oocyte complexes in vitro: a multistep analysis. Amino Acids, 2024. 56(1): p. 2.

94. Feng, X., et al., Heat-Stress Impacts on Developing Bovine Oocytes: Unraveling Epigenetic Changes, Oxidative Stress, and Developmental Resilience. Int J Mol Sci, 2024. 25(9).

95. Ju, J.C. and J.K. Tseng, Nuclear and cytoskeletal alterations of in vitro matured porcine oocytes under hyperthermia. Mol Reprod Dev, 2004. 68(1): p. 125–33.

96. Tseng, J.K., et al., Influences of follicular size on parthenogenetic activation and in vitro heat shock on the cytoskeleton in cattle oocytes. Reprod Domest Anim, 2004. 39(3): p. 146–53.

97. Ascari, I.J., et al., Addition of insulin-like growth factor I to the maturation medium of bovine oocytes subjected to heat shock: effects on the production of reactive oxygen species, mitochondrial activity and oocyte competence. Domest Anim Endocrinol, 2017. 60: p. 50–60.

98. Soto, P. and L.C. Smith, BH4 peptide derived from Bcl-xL and Bax-inhibitor peptide suppresses apoptotic mitochondrial changes in heat stressed bovine oocytes. Mol Reprod Dev, 2009. 76(7): p. 637–46.

99. Amaral, C.S., et al., Heat stress on oocyte or zygote compromises embryo development, impairs interferon tau production and increases reactive oxygen species and oxidative stress in bovine embryos produced in vitro. Mol Reprod Dev, 2020. 87(8): p. 899–909.

100. Camargo, L.S.A., et al., Contrasting effects of heat shock during in vitro maturation on development of in vitro-fertilized and parthenogenetic bovine embryos. Reprod Domest Anim, 2019. 54(10): p. 1357–1365.

101. Cavallari, F.C., et al., Effects of melatonin on production of reactive oxygen species and developmental competence of bovine oocytes exposed to heat shock and oxidative stress during in vitro maturation. Zygote, 2019. 27(3): p. 180–186.

102. Edwards, J.L., et al., Developmental competence of bovine embryos from heat-stressed ova. J Dairy Sci, 2009. 92(2): p. 563–70.

103. Edwards, J.L. and P.J. Hansen, Elevated temperature increases heat shock protein 70 synthesis in bovine two-cell embryos and compromises function of maturing oocytes. Biol Reprod, 1996. 55(2): p. 341–6.

104. Ju, J.C., J.E. Parks, and X. Yang, Thermotolerance of IVM-derived bovine oocytes and embryos after short-term heat shock. Mol Reprod Dev, 1999. 53(3): p. 336–40.

105. Meiyu, Q., D. Liu, and Z. Roth, IGF-I slightly improves nuclear maturation and cleavage rate of bovine oocytes exposed to acute heat shock in vitro. Zygote, 2015. 23(4): p. 514–24.

106. Payton, R.R., et al., Mitochondrial-related consequences of heat stress exposure during bovine oocyte maturation persist in early embryo development. J Reprod Dev, 2018. 64(3): p. 243–251.

107. Pohland, R., et al., Influence of long-term thermal stress on the in vitro maturation on embryo development and Heat Shock Protein abundance in zebu cattle. Anim Reprod, 2020. 17(3): p. e20190085.

108. Rodrigues, T.A., et al., Follicular fluid exosomes act on the bovine oocyte to improve oocyte competence to support development and survival to heat shock. Reprod Fertil Dev, 2019. 31(5): p. 888–897.

109. Roth, Z. and P.J. Hansen, Sphingosine 1-phosphate protects bovine oocytes from heat shock during maturation. Biol Reprod, 2004. 71(6): p. 2072–8.

110. Roth, Z. and P.J. Hansen, Involvement of apoptosis in disruption of developmental competence of bovine oocytes by heat shock during maturation. Biol Reprod, 2004. 71(6): p. 1898–906.

111. Souza, V., et al., Heat shock during in vitro maturation of bovine oocytes disturbs bta-miR-19b and DROSHA transcripts abundance after in vitro fertilization. Reprod Domest Anim, 2021. 56(8): p. 1128–1136.

112. Stamperna, K., et al., Developmental competence of heat stressed oocytes from Holstein and Limousine cows matured in vitro. Reprod Domest Anim, 2021. 56(10): p. 1302–1314.

113. Wrzecinska, M., et al., Disorder of Biological Quality and Autophagy Process in Bovine Oocytes Exposed to Heat Stress and the Effectiveness of In Vitro Fertilization. Int J Mol Sci, 2023. **24**(13).

114. Schrock, G.E., et al., Early in vitro fertilization improves development of bovine ova heat stressed during in vitro maturation. J Dairy Sci, 2007. 90(9): p. 4297–303.

115. Payton, R.R., et al., Impact of heat stress exposure during meiotic maturation on oocyte, surrounding cumulus cell, and embryo RNA populations. J Reprod Dev, 2011. 57(4): p. 481–91.

116. Balboula, A.Z., et al., Cathepsin B activity has a crucial role in the developmental competence of bovine cumulus-oocyte complexes exposed to heat shock during in vitro maturation. Reproduction, 2013. 146(4): p. 407–17.

117. Current, J.Z., M. Mentler, and B.D. Whitaker, Linoleic and linolenic acids reduce the effects of heat stress-induced damage in pig oocytes during maturation in vitro. In Vitro Cell Dev Biol Anim, 2022. 58(7): p. 599–609.

118. Lee, J., et al., Effects of heat stress on conception in Holstein and Jersey cattle and oocyte maturation in vitro. J Anim Sci Technol, 2023. 65(2): p. 324–335.

119. Stamperna, K., et al., Short term temperature elevation during IVM affects embryo yield and alters gene expression pattern in oocytes, cumulus cells and blastocysts in cattle. Theriogenology, 2020. 156: p. 36–45.

120. Duncan, F.E., R. Confino, and M.E. Pavone, Chapter 9 - Female Reproductive Aging: From Consequences to Mechanisms, Markers, and Treatments, in Conn’s Handbook of Models for Human Aging (Second Edition), J.L. Ram and P.M. Conn, Editors. 2018, Academic Press. p. 109-130.

121. Faheem, M.S., et al., Adaptive and Biological Responses of Buffalo Granulosa Cells Exposed to Heat Stress under In Vitro Condition. Animals (Basel), 2021. **11**(3).

122. Li, L., et al., The effect of heat stress on gene expression, synthesis of steroids, and apoptosis in bovine granulosa cells. Cell Stress Chaperones, 2016. 21(3): p. 467–75.

123. Klabnik, J.L., et al., Heat-induced increases in body temperature in lactating dairy cows: impact on the cumulus and granulosa cell transcriptome of the periovulatory follicle. Journal of Animal Science, 2022. 100(7).

124. Karaşahin, T., Effect of low doses of FSH and season on the in vitro maturation, fertilization and embryo development of bovine oocytes. The Indian journal of animal sciences, 2020. 90: p. 564–568.

125. Toosinia, S., et al., Ameliorating Effect of Sodium Selenite on Developmental and Molecular Response of Bovine Cumulus-Oocyte Complexes Matured in Vitro Under Heat Stress Condition. Biol Trace Elem Res, 2024. 202(1): p. 161–174.

126. Shi, X.Y., et al., Idebenone relieves the damage of heat stress on the maturation and developmental competence of porcine oocytes. Reprod Domest Anim, 2022. 57(4): p. 418–428.

127. Peng, K., et al., Mogroside V alleviates the heat stress-induced disruption of the porcine oocyte in vitro maturation. Theriogenology, 2024. 217: p. 37–50.

128. Sammad, A., et al., Vitamin C Alleviates the Negative Effects of Heat Stress on Reproductive Processes by Regulating Amino Acid Metabolism in Granulosa Cells. Antioxidants (Basel), 2024. 13(6).

