## Supplemental Table 1 for "The Effects of Heat Stress on the Ovary, Follicles and Oocytes: A Systematic Review"

**Supplementary Table SI**. Search strategy across the databases

| **PubMed** | | |
| --- | --- | --- |
|  | Search Strategy | Publications |
| #1 | (embryo* [tiab] OR Follicle*[tiab] OR follicular*[tiab] OR oocyte*[tiab] OR ovum[tiab] OR ovar*[tiab] OR ovulat*[tiab] OR Ovarian-reserve*[tiab] OR Anti-mullerian-hormone*[tiab] OR AMH[tiab] OR Antral-follicle-count*[tiab] OR Estradiol*[tiab] OR Follicle-stimulating-hormone*[tiab] OR FSH[tiab] OR egg[tiab] OR eggs[tiab] OR gamete*[tiab] OR reproductive-health*[tiab] OR "Embryonic Structures"[Mesh] OR "Ovarian Follicle"[Mesh] OR "Oocytes"[Mesh] OR "Ovum"[Mesh] OR "Ovary"[Mesh] OR "Ovulation"[Mesh] OR "Ovarian Reserve"[Mesh] OR "Anti-Mullerian Hormone"[Mesh] OR "Estradiol"[Mesh] OR "Follicle Stimulating Hormone"[Mesh] OR "Germ Cells"[Mesh] OR "Reproductive Health"[Mesh]) | 1446792 |
| #2 | (Pregnancy[MeSH] OR "Birth weight"[MeSH] OR "pregnancy outcome"[MeSH] OR "premature birth"[MeSH] OR "infant, low birth weight"[MeSH] OR "pregnancy complications"[MeSH] OR "gestational age"[MeSH] OR "Postpartum Hemorrhage"[Mesh] OR "infant, premature"[MeSH] OR "Maternal Death"[Mesh] OR "perinatal death"[MeSH] OR "Intensive Care Units, Neonatal"[Mesh] OR "Obstetric Labor Complications"[Mesh] OR "maternal mortality"[MeSH] OR "fetal death"[MeSH] OR Fertility[MeSH] OR infertility[MeSH] OR "Fertilization"[Mesh] OR "Birth Rate"[Mesh] OR "Reproductive Health"[Mesh] OR pregnan*[tiab] OR conception[tiab] OR Birth-rate*[tiab] OR birth-weight[tiab] OR birth-outcome*[tiab] OR preterm-delivery[tiab] OR LBW[tiab] OR premature-birth*[tiab] OR adverse-perinatal-outcome*[tiab] OR pregnancy-complication*[tiab] OR preterm-birth*[tiab] OR premature-infant*[tiab] OR gestational-age[tiab] OR Stillbirth[tiab] OR postpartum-hemorrhage*[tiab] OR matern*[tiab] OR maternal-death[tiab] OR maternal-ICU[tiab] OR perinatal-death*[tiab] OR neonatal-death*[tiab] OR NICU[tiab] OR neonatal-ICU[tiab] OR maternal-mortality[tiab] OR perinatal-outcome*[tiab] OR obstetric-labor-complication*[tiab] OR Fetal-death[tiab] OR fetal-demise[tiab] OR severe-maternal-morbidity[tiab] OR Fertil*[tiab] OR Reproductive-health[tiab] OR female-reproduction[tiab] OR Female-fertil*[tiab] OR Infertil*[tiab] OR Subfertil*[tiab] OR (("Delivery, Obstetric"[Mesh] OR "Obstetrics"[Mesh] OR delivery[tiab] OR obstetric[tiab] OR neonatal[tiab] OR perinatal[tiab] OR antenatal[tiab]) AND (outcome*[tiab]))) | 1727604 |
| #3 | ("global warming"[MeSH Terms] OR "Climate Change"[Mesh:NoExp] OR "Heat-Shock Response"[Mesh] OR "Extreme Heat"[Mesh] OR "global warming"[Title/Abstract] OR "hot temperature*"[Title/Abstract] OR "high temperature*"[Title/Abstract] OR "hot environment*"[Title/Abstract] OR "heat stress*"[Title/Abstract] OR "environmental monitor*"[Title/Abstract] OR "extreme temperature*"[Title/Abstract] OR "extreme heat*"[Title/Abstract] OR "heat wave*"[Title/Abstract] OR climate-change[tiab] OR climate-data[tiab] OR ambient-temperature*[tiab]) | 193667 |
| #4 | #1 AND #2 AND #3 | 1641 |

| **Embase** | | |
| --- | --- | --- |
|  | Search strategy | Publications |
| #1 | ('embryo'/exp OR 'oocyte'/exp OR 'ovum'/exp OR 'ovarian reserve'/exp OR 'Muellerian inhibiting factor'/exp OR 'antral follicle count'/exp OR 'estradiol'/exp OR 'follitropin'/exp OR 'gamete'/exp OR 'reproductive health'/exp OR embryo*:ab,ti OR Follicle*:ab,ti OR follicular*:ab,ti OR oocyte*:ab,ti OR ovum:ab,ti OR ovar*:ab,ti OR ovulat*:ab,ti OR Ovarian-reserve*:ab,ti OR Anti-mullerian-hormone*:ab,ti OR AMH:ab,ti OR Antral-follicle-count*:ab,ti OR Estradiol*:ab,ti OR Follicle-stimulating-hormone*:ab,ti OR FSH:ab,ti OR egg:ab,ti OR eggs:ab,ti OR gamete*:ab,ti OR reproductive-health*:ab,ti) | 1530669 |
| #2 | (('pregnancy'/exp OR 'birth weight'/exp OR 'pregnancy outcome'/exp OR 'prematurity'/exp OR 'low birth weight'/exp OR 'pregnancy complication'/exp OR 'gestational age'/exp OR 'postpartum hemorrhage'/exp OR 'maternal death'/exp OR 'perinatal death'/exp OR 'neonatal intensive care unit'/exp OR 'maternal mortality'/exp OR 'fetus death'/exp OR 'fertility'/exp OR 'infertility'/exp OR 'conception'/exp OR 'birth rate'/exp OR 'reproductive health'/exp OR 'premature labor'/exp OR 'stillbirth'/exp OR 'newborn death'/exp OR 'female fertility'/exp OR pregnan*:ab,ti OR conception:ab,ti OR Birth-rate*:ab,ti OR birth-weight:ab,ti OR birth-outcome*:ab,ti OR preterm-delivery:ab,ti OR LBW:ab,ti OR premature-birth*:ab,ti OR adverse-perinatal-outcome*:ab,ti OR pregnancy-complication*:ab,ti OR preterm-birth*:ab,ti OR premature-infant*:ab,ti OR gestational-age:ab,ti OR Stillbirth:ab,ti OR postpartum-hemorrhage*:ab,ti OR matern*:ab,ti OR maternal-death:ab,ti OR maternal-ICU:ab,ti OR perinatal-death*:ab,ti OR neonatal-death*:ab,ti OR NICU:ab,ti OR neonatal-ICU:ab,ti OR maternal-mortality:ab,ti OR perinatal-outcome*:ab,ti OR obstetric-labor-complication*:ab,ti OR Fetal-death:ab,ti OR fetal-demise:ab,ti OR severe-maternal-morbidity:ab,ti OR Fertil*:ab,ti OR Reproductive-health:ab,ti OR female-reproduction:ab,ti OR Female-fertil*:ab,ti OR Infertil*:ab,ti OR Subfertil*:ab,ti) OR (('obstetric delivery'/exp OR 'obstetrics'/exp OR delivery:ab,ti OR obstetric:ab,ti OR neonatal:ab,ti OR perinatal:ab,ti OR antenatal:ab,ti) AND (outcome*:ab,ti))) | 2182173 |
| #3 | ('greenhouse effect'/exp OR 'climate change'/exp OR 'heat stress'/exp OR 'extreme hot weather'/exp OR 'heat wave'/exp OR "global warming":ab,ti OR "hot temperature*":ab,ti OR "high temperature*":ab,ti OR "hot environment*":ab,ti OR "heat stress*":ab,ti OR "environmental monitor*":ab,ti OR "extreme temperature*":ab,ti OR "extreme heat*":ab,ti OR "heat wave*":ab,ti OR climate-change:ab,ti OR climate-data:ab,ti OR ambient-temperature*:ab,ti) | 213834 |
| #4 | #1 AND # 2 AND #3 | 1686 |

| **Cochrane** | | |
| --- | --- | --- |
|  | Search strategy | Publications |
| #1 | MeSH descriptor: [Embryonic Structures] explode all trees | 4248 |
| #2 | MeSH descriptor: [Ovarian Follicle] explode all trees | 647 |
| #3 | MeSH descriptor: [Oocytes] explode all trees | 572 |
| #4 | MeSH descriptor: [Ovum] explode all trees | 894 |
| #5 | MeSH descriptor: [Ovary] explode all trees | 1438 |
| #6 | MeSH descriptor: [Ovulation] explode all trees | 950 |
| #7 | MeSH descriptor: [Ovarian Reserve] explode all trees | 144 |
| #8 | MeSH descriptor: [Anti-Mullerian Hormone] explode all trees | 161 |
| #9 | MeSH descriptor: [Estradiol] explode all trees | 5013 |
| #10 | MeSH descriptor: [Follicle Stimulating Hormone] explode all trees | 2187 |
| #11 | MeSH descriptor: [Germ Cells] explode all trees | 1349 |
| #12 | MeSH descriptor: [Reproductive Health] explode all trees | 221 |
| #13 | (embryo* OR Follicle* OR follicular* OR oocyte* OR ovum OR ovar* OR ovulat* OR Ovarian-reserve* OR Anti-mullerian-hormone* OR AMH OR Antral-follicle-count* OR Estradiol* OR Follicle-stimulating-hormone* OR FSH OR egg OR eggs OR gamete* OR reproductive-health*):ti,ab,kw | 52179 |
| #14 | #1 OR #2 OR #4 OR #6 OR #7 OR #8 OR #9 OR #10 OR #11 OR #12 OR #13 | 55951 |
| #15 | MeSH descriptor: [Pregnancy] explode all trees | 31569 |
| #16 | MeSH descriptor: [Birth Weight] explode all trees | 2563 |
| #17 | MeSH descriptor: [Pregnancy Outcome] explode all trees | 5164 |
| #18 | MeSH descriptor: [Premature Birth] explode all trees | 2162 |
| #19 | MeSH descriptor: [Infant, Low Birth Weight] explode all trees | 2658 |
| #20 | MeSH descriptor: [Pregnancy Complications] explode all trees | 16219 |
| #21 | MeSH descriptor: [Gestational Age] explode all trees | 3985 |
| #22 | MeSH descriptor: [Postpartum Hemorrhage] explode all trees | 985 |
| #23 | MeSH descriptor: [Infant, Premature] explode all trees | 4956 |
| #24 | MeSH descriptor: [Maternal Death] explode all trees | 51 |
| #25 | MeSH descriptor: [Perinatal Death] explode all trees | 165 |
| #26 | MeSH descriptor: [Intensive Care Units, Neonatal] explode all trees | 1037 |
| #27 | MeSH descriptor: [Obstetric Labor Complications] explode all trees | 5333 |
| #28 | MeSH descriptor: [Maternal Mortality] explode all trees | 216 |
| #29 | MeSH descriptor: [Fetal Death] explode all trees | 547 |
| #30 | MeSH descriptor: [Fertility] explode all trees | 639 |
| #31 | MeSH descriptor: [Infertility] explode all trees | 4494 |
| #32 | MeSH descriptor: [Fertilization] explode all trees | 374 |
| #33 | MeSH descriptor: [Birth Rate] explode all trees | 311 |
| #34 | MeSH descriptor: [Reproductive Health] explode all trees | 221 |
| #35 | MeSH descriptor: [Delivery, Obstetric] explode all trees | 7288 |
| #36 | MeSH descriptor: [Obstetrics] explode all trees | 494 |
| #37 | (delivery OR obstetric OR neonatal OR perinatal OR antenatal):ti,ab,kw | 83499 |
| #38 | (outcome*):ti,ab,kw | 777559 |
| #39 | (pregnan* OR conception OR Birth-rate* OR birth-weight OR birth-outcome* OR preterm-delivery OR LBW OR premature-birth* OR adverse-perinatal-outcome* OR pregnancy-complication* OR preterm-birth* OR premature-infant* OR gestational-age OR Stillbirth OR postpartum-hemorrhage* OR matern* OR maternal-death OR maternal-ICU OR perinatal-death* OR neonatal-death* OR NICU OR neonatal-ICU OR maternal-mortality OR perinatal-outcome* OR obstetric-labor-complication* OR Fetal-death OR fetal-demise OR severe-maternal-morbidity OR Fertil* OR Reproductive-health OR female-reproduction OR Female-fertil* OR Infertil* OR Subfertil*):ti,ab,kw | 118590 |
| #40 | #15 OR #16 OR #17 OR #18 OR #19 OR #20 OR #21 OR #22 OR #23 OR #24 OR #25 OR #26 OR #27 OR #28 OR #29 OR #30 OR #31 OR #32 OR #33 OR #34 OR #39 OR ((#35 OR #36 OR #37) AND #38) | 141566 |
| #41 | MeSH descriptor: [Global Warming] explode all trees | 4 |
| #42 | MeSH descriptor: [Climate Change] this term only | 38 |
| #43 | MeSH descriptor: [Heat-Shock Response] explode all trees | 116 |
| #44 | MeSH descriptor: [Extreme Heat] explode all trees | 9 |
| #45 | (global-warming OR hot-temperature* OR high-temperature* OR hot-environment* OR heat-stress* OR environmental-monitor* OR extreme-temperature* OR extreme-heat* OR heat-wave* OR climate-change OR climate-data OR ambient-temperature*):ti,ab,kw | 4316 |
| #46 | #41 OR #42 OR #43 OR #44 OR #45 | 4329 |
| #47 | #14 AND #40 AND #46 | 46 |

**1 Cochrane review; 45 trials*

| **Scopus** | | |
| --- | --- | --- |
|  | Search strategy | Publications |
| #1 | TITLE-ABS ( embryo* OR follicle* OR follicular* OR oocyte* OR ovum OR ovar* OR ovulat* OR ovarian-reserve* OR anti-mullerian-hormone* OR amh OR antral-follicle-count* OR estradiol* OR follicle-stimulating-hormone* OR fsh OR egg OR eggs OR gamete* OR reproductive-health* ) | 1393008 |
| #2 | TITLE-ABS ( pregnan* OR conception OR birth-rate* OR birth-weight OR birth-outcome* OR preterm-delivery OR lbw OR premature-birth* OR low-birth-weight OR adverse-perinatal-outcome* OR pregnancy-complication* OR preterm-birth* OR premature-infant* OR gestational-age OR stillbirth OR postpartum-hemorrhage* OR matern* OR maternal-death OR maternal-icu OR perinatal-death* OR neonatal-death* OR neonatal-intensive-care-unit* OR nicu OR neonatal-icu OR maternal-mortality OR perinatal-outcome* OR obstetric-labor-complication* OR fetal-death OR fetal-demise OR severe-maternal-morbidity OR fertil* OR reproductive-health OR female-reproduction OR female-fertil* OR infertil* OR subfertil* OR ( ( delivery OR obstetric OR neonatal OR perinatal OR antenatal ) AND ( outcome* ) ) ) | 1893301 |
| #3 | TITLE-ABS ( global-warming OR hot-temperature* OR high-temperature* OR hot-environment* OR heat-stress* OR environmental-monitor* OR extreme-temperature* OR extreme-heat* OR heat-wave* OR climate-change OR climate-data OR ambient-temperature* ) | 1181890 |
| #4 | #1 AND #2 AND #3 | 1793 |

| **Global Health** | | |
| --- | --- | --- |
|  | Search strategy | Publications |
| #1 | (DE "embryos" OR DE "oocytes" OR DE "ova" OR DE "ovulation" OR DE "estradiol” OR DE "FSH" OR DE "gametes" OR TI (embryo* OR Follicle* OR follicular* OR oocyte* OR ovum OR ovar* OR ovulat* OR Ovarian-reserve* OR Anti-mullerian-hormone* OR AMH OR Antral-follicle-count* OR Estradiol* OR Follicle-stimulating-hormone* OR FSH OR egg OR eggs OR gamete* OR reproductive-health*) OR AB (embryo* OR Follicle* OR follicular* OR oocyte* OR ovum OR ovar* OR ovulat* OR Ovarian-reserve* OR Anti-mullerian-hormone* OR AMH OR Antral-follicle-count* OR Estradiol* OR Follicle-stimulating-hormone* OR FSH OR egg OR eggs OR gamete* OR reproductive-health*)) | 120998 |
| #2 | (DE "pregnancy" OR DE "birth weight" OR DE "low birth weight infants" OR DE "pregnancy complications" OR DE "premature infants" OR DE "maternal mortality" OR DE "perinatal mortality" OR DE "fetal death" OR DE "fertility" OR DE "infertility" OR DE "conception" OR DE "birth rate" OR TI ( pregnan* OR conception OR Birth-rate* OR birth-weight OR birth-outcome* OR preterm-delivery OR LBW OR premature-birth* OR adverse-perinatal-outcome* OR pregnancy-complication* OR preterm-birth* OR premature-infant* OR gestational-age OR Stillbirth OR postpartum-hemorrhage* OR matern* OR maternal-death OR maternal-ICU OR perinatal-death* OR neonatal-death* OR NICU OR neonatal-ICU OR maternal-mortality OR perinatal-outcome* OR obstetric-labor-complication* OR Fetal-death OR fetal-demise OR severe-maternal-morbidity OR Fertil* OR Reproductive-health OR female-reproduction OR Female-fertil* OR Infertil* OR Subfertil* ) OR AB ( pregnan* OR conception OR Birth-rate* OR birth-weight OR birth-outcome* OR preterm-delivery OR LBW OR premature-birth* OR adverse-perinatal-outcome* OR pregnancy-complication* OR preterm-birth* OR premature-infant* OR gestational-age OR Stillbirth OR postpartum-hemorrhage* OR matern* OR maternal-death OR maternal-ICU OR perinatal-death* OR neonatal-death* OR NICU OR neonatal-ICU OR maternal-mortality OR perinatal-outcome* OR obstetric-labor-complication* OR Fetal-death OR fetal-demise OR severe-maternal-morbidity OR Fertil* OR Reproductive-health OR female-reproduction OR Female-fertil* OR Infertil* OR Subfertil* ) OR ((TI ( delivery OR obstetric OR neonatal OR perinatal OR antenatal ) OR AB ( delivery OR obstetric OR neonatal OR perinatal OR antenatal) OR DE "obstetrics") AND (TI outcome* OR AB outcome*)) | 255675 |
| #3 | (DE "climatic change" OR TI (global-warming OR hot-temperature* OR high-temperature* OR hot-environment* OR heat-stress* OR environmental-monitor* OR extreme-temperature* OR extreme-heat* OR heat-wave* OR climate-change OR climate-data OR ambient-temperature*) OR AB (global-warming OR hot-temperature* OR high-temperature* OR hot-environment* OR heat-stress* OR environmental-monitor* OR extreme-temperature* OR extreme-heat* OR heat-wave* OR climate-change OR climate-data OR ambient-temperature*) ) | 33524 |
| #4 | #1 AND #2 AND #3 | 120 |
